# High-resolution structure and dynamics of mitochondrial complex I – insights into the proton pumping mechanism

**DOI:** 10.1101/2021.04.16.440187

**Authors:** Kristian Parey, Jonathan Lasham, Deryck J. Mills, Amina Djurabekova, Outi Haapanen, Etienne Galemou Yoga, Hao Xie, Werner Kühlbrandt, Vivek Sharma, Janet Vonck, Volker Zickermann

## Abstract

Mitochondrial NADH:ubiquinone oxidoreductase (complex I) is a 1 MDa membrane protein complex with a central role in energy metabolism. Redox-driven proton translocation by complex I contributes substantially to the proton motive force that drives ATP synthase. Several structures of complex I from bacteria and mitochondria have been determined but its catalytic mechanism has remained controversial. We here present the cryo-EM structure of complex I from *Yarrowia lipolytica* at 2.1 Å resolution, which reveals the positions of more than 1600 protein-bound water molecules, of which ∼100 are located in putative proton translocation pathways. Another structure of the same complex under steady-state activity conditions at 3.4 Å resolution indicates conformational transitions that we associate with proton injection into the central hydrophilic axis. By combining high-resolution structural data with site-directed mutagenesis and large-scale molecular dynamics simulations, we define details of the proton translocation pathways, and offer new insights into the redox-coupled proton pumping mechanism of complex I.

## Introduction

Mitochondrial NADH:ubiquinone oxidoreductase (complex I) is the largest and most intricate membrane protein complex of the respiratory chain (Galemou Yoga et al., 2020a; Haapanen and Sharma, 2018; Hirst, 2013; Sazanov, 2015). Complex I couples the transfer of electrons from NADH to ubiquinone with the translocation of protons across the inner mitochondrial membrane, with a stoichiometry of 4 H^+^ per NADH (Galkin et al., 2006; Galkin et al., 1999; Jones et al., 2017; Wikström, 1984). Redox-linked proton translocation by complex I contributes substantially to the proton motive force that drives ATP synthase. Complex I dysfunction is associated with neuromuscular and neurodegenerative diseases (Fiedorczuk and Sazanov, 2018; Rodenburg, 2016). The structure of complex I has been determined by X-ray crystallography (Baradaran et al., 2013; Zickermann et al., 2015) and cryo-EM at increasing resolution (Agip et al., 2019; Grba and Hirst, 2020; Kampjut and Sazanov, 2020; Klusch et al., 2021; Parey et al., 2019; Parey et al., 2020; Soufari et al., 2020), but the molecular details of the coupling mechanism are not fully understood and remain controversial.

The L-shaped enzyme complex has three functional modules. The NADH oxidation module (N module) and ubiquinone (Q) reduction module (Q module) together constitute the peripheral arm. The membrane arm has a proximal and a distal proton translocation module (P_P_ and P_D_ module). The Q module is located above the membrane plane so that the hydrophobic quinone substrate has to pass through a ∼35 Å tunnel to reach the Q reduction site near FeS cluster N2. In the P module, proton translocation pathways from the negative (N) side to the positive (P) side of the membrane were assigned to subunits ND2, ND4 and ND5 that resemble Mrp-type sodium-proton antiporter subunits (Mathiesen and Hägerhäll, 2002). A fourth proton channel has long been anticipated but its position remains unknown. It is generally accepted that Q reduction releases the energy for proton translocation, but the proton pumping sites are up to 200 Å away from the Q reduction site. Several lines of evidence indicate that a set of conserved loops at the interface between the Q and P_P_ modules plays a key role in energy conversion (Cabrera-Orefice et al., 2018; Galemou Yoga et al., 2019; Galemou Yoga et al., 2020b; Grba and Hirst, 2020; Kampjut and Sazanov, 2020; Zickermann et al., 2015). The loop connecting transmembrane helices (TMHs) 5 and 6 of membrane subunit ND1 (the ND1 loop) harbors a cluster of strictly conserved acidic residues close to the Q reduction site and lines part of the Q tunnel. A chain of charged residues in the membrane-embedded central subunits forms an extended hydrophilic axis that connects ND1 with the proton pumping sites. Recent cryo-EM structures at 2.7 Å (Grba and Hirst, 2020) and 2.5-2.9 Å (Kampjut and Sazanov, 2020) resolution indicated water molecules that define putative proton translocation pathways in the P module.

The dynamics of complex I have been studied by molecular simulations (Haapanen and Sharma, 2018; Hummer and Wikström, 2016; Kaila, 2018) and by determining its structure in different functional states (Agip et al., 2018; Gutierrez-Fernandez et al., 2020; Kampjut and Sazanov, 2020; Parey et al., 2018). For mammalian and plant complex I, a change of the relative orientation of peripheral arm and membrane arm was found in conjunction with well-defined changes in several central subunits (Agip et al., 2018; Kampjut and Sazanov, 2020; Klusch et al., 2021; Letts et al., 2019). We have previously analyzed substrate binding to complex I during steady state activity (Parey et al., 2018) but at the time, the resolution of our cryo-EM structure did not allow us to identify molecular changes in detail.

As a characteristic feature, complex I undergoes a reversible transition between an active (A) and a deactive (D) form (Kotlyar and Vinogradov, 1990). The A form converts into the D form when complex I is idle in the absence of substrates. The reconversion into the A form takes several turnover cycles. Deactivation of complex I is thought to protect against excessive ROS formation during ischemia/ reperfusion (Chouchani et al., 2013), but the structural basis of the A/D transition is not clear. Biochemical analysis has shown that complex I purified from *Y. lipolytica* is in the D form (Parey et al., 2019) as a default ground state. The energy barrier between A and D form in fungal complex I is smaller (Maklashina et al., 2003) and deactivation appears to be less stringent than in the mammalian complex, because structural hallmarks of the D form described for mammalian complex I, including the pronounced weakening of loop densities in and around the Q binding site (Agip et al., 2018) and the striking relocation of TMH4 in ND6 (Kampjut and Sazanov, 2020) are not observed.

Here, we present the 2.1 Å resolution structure of respiratory complex I from the yeast *Yarrowia lipolytica* in the D form and a 3.4 Å resolution structure of the same complex captured under turnover conditions. Our data provide a comprehensive picture of protein-bound water molecules in the P module. Under turnover conditions, we observe structural changes in the ND1 subunit that forms the link between Q and P modules. Large-scale atomistic molecular dynamics (MD) simulations based on our high-resolution structure reveal how the protonation states of key amino acid residues control the hydration and dynamics of regions that are crucial for the coupling mechanism.

### High-resolution structure

Respiratory complex I from the aerobic yeast *Y. lipolytica* in the D form was purified in the detergent lauryl maltose neopentyl glycol (LMNG) by His-tag affinity and size exclusion chromatography (Parey et al., 2019). To optimize the preparation for cryo-EM we added an extra purification step by ion exchange chromatography. The sample had a high Q reductase activity of more than 20 μmol·min^−1^·mg^−1^. Note that we measure this activity without reactivation by lipids in the assay buffer, indicating that our preparation is catalytically fully competent when applied to the cryo-EM grid. Single-particle analysis yielded a map with an average resolution of 2.4 Å (Figure S1). Density modification (Terwilliger et al., 2020) improved the overall resolution to 2.1 Å, especially of some less-ordered regions and many water molecules (Figure 1, Figure S2). The core of the peripheral arm and membrane arm are particularly well-resolved, as indicated by the well-separated iron and sulfur atoms in the FeS clusters and low central density of aromatic side chains (Figure 1, Figure S2, Movie 1). At this resolution, the main-chain carbonyl oxygens are clearly visible and the rotamers of many side chains can be distinguished (Figure 1, Figure S2). Our model contains 8035 residues, all cofactors, 34 lipid molecules, 2 LMNG detergent molecules and 1617 water molecules. Further, N-formyl methionines were visible in several of the mitochondrially-encoded subunits (Figure S2), as was the dimethyl arginine residue 121 of NDUFS2 (Figure 1). The density for the native Q9 molecule observed in the Q tunnel (Parey et al., 2019) appeared weaker, most likely due to its partial removal during the additional purification step. In a *Y. lipolytica* complex I purified in dodecyl maltoside (DDM) (Grba and Hirst, 2020), this site had been occupied by a detergent molecule. In our present structure the same site is essentially empty, suggesting that the bulkier LMNG cannot enter the narrow substrate tunnel. All previously observed lipid molecules (Parey et al., 2019) were retained, indicating that they are tightly bound and therefore an integral part of the complex. Interestingly, depletion of Q in the tunnel binding site is linked with a loss of density for a five-residue segment of the functionally important loop connecting TMH1 and 2 of ND3 (Cabrera-Orefice et al., 2018), suggesting a functional connection between Q binding and the mobility of this loop (Haapanen et al., 2020).

**Figure 1.**
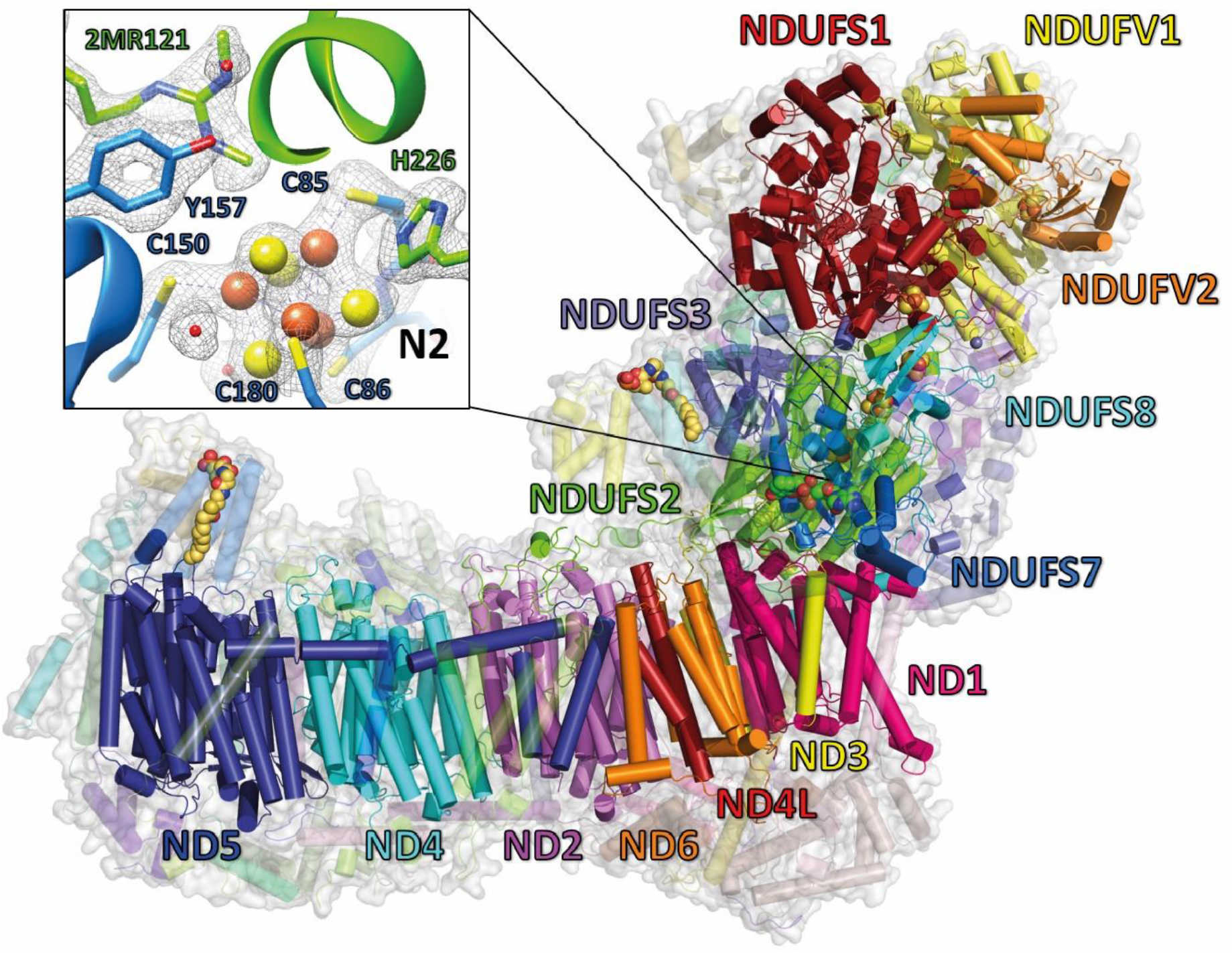
Overview of central subunits and detailed view on FeS cluster N2. Side view of *Y. lipolytica* complex I. The 14 central subunits are labeled and shown in color, accessory subunits are transparent. The inset shows the environment of FeS cluster N2 with the cryo-EM density (grey mesh). See also Figure S1 and S2 and Table S1.

### Internal volumes and protein-bound water molecules in the membrane arm

We used the software tool Caver (Chovancova et al., 2012) to identify tunnels and cavities in the ND1 subunit. In agreement with previous studies (Grba and Hirst, 2020; Kampjut and Sazanov, 2020), we found that this subunit encloses a substantial branched cavity (Figure 2). The main branches are (i) the access channel for Q towards the active site in subunits NDUFS2 and NDUFS7, and (ii) a ∼25 Å tunnel extending towards the interface with subunit ND6 in the membrane arm. The tunnel leading to the membrane interior is referred to as the E channel (Baradaran et al., 2013) because it is lined by the strictly conserved and functionally important glutamates Glu196^ND1^, Glu231^ND1^ and Glu147^ND1^ (Kurki et al., 2000). One water molecule (W6) was modelled at the entrance and five water molecules close to the distal end of the E channel (W8-W12) (Figure 2, Figure S3). Four water molecules (W19-W22) are bound at the junction of subunits ND1, ND3 and ND6. They interact with Ser114, Thr151 and Glu147 of ND1 and with the strictly conserved Asp67 of ND3. TMH3 of ND6 is a π-helix with two water molecules (W20, W22) bound by backbone interactions in the π-bulge region. An ∼18 Å dry region separates the waters in ND1 from an extended series of water molecules that originates at the interface of subunits ND6 and ND4L (Figure 2). The strictly conserved glutamates of ND4L, Glu30 and Glu66, and the neighbouring Glu131 of ND2 are highly hydrated (W25-W31). One water molecule (W23) is ligated by Tyr59^ND6^ and the highly conserved Tyr63^ND6^ of the ND6 π-bulge helix. The adjacent ND2 subunit comprises a central water cluster and is traversed horizontally by a 44 Å chain of water molecules. Remarkably, all gaps in this chain are filled by hydrophilic or charged residues, most of which are strictly conserved (Figure 2). In contrast to previous reports (Grba and Hirst, 2020; Kampjut and Sazanov, 2020), an extension of the central water cluster at Lys241^ND2^ towards Lys282^ND2^ and Arg283^ND2^ indicates a clear entry point for protons at the N-side end of TMH10 (Figure 2). Interestingly, the Lys in TMH10 is also conserved in ND4 and the Lys-Arg motif is conserved in TMH10 of ND5 (Figure S4), suggesting similar N-side entry portals for protons for all three subunits. Intriguingly, fewer water molecules were detected in subunits ND4 and ND5 than in ND2. Overall, the pattern of water molecules in ND4 resembles that in ND2, but gaps between water molecules or clusters are larger. Water molecules are found near the N-side TMH10 Lys, but the pathway into the subunit interior is less clear. In ND5, which differs from ND2 and ND4 at the sequence level (Figure S4) (Djurabekova et al., 2020), water molecules line up along the strictly conserved ND5 residues Glu144, Arg175, Asp178 and Lys226, while single water molecules reside close to conserved histidines (His251^ND5^ and His335^ND5^).

**Figure 2:**
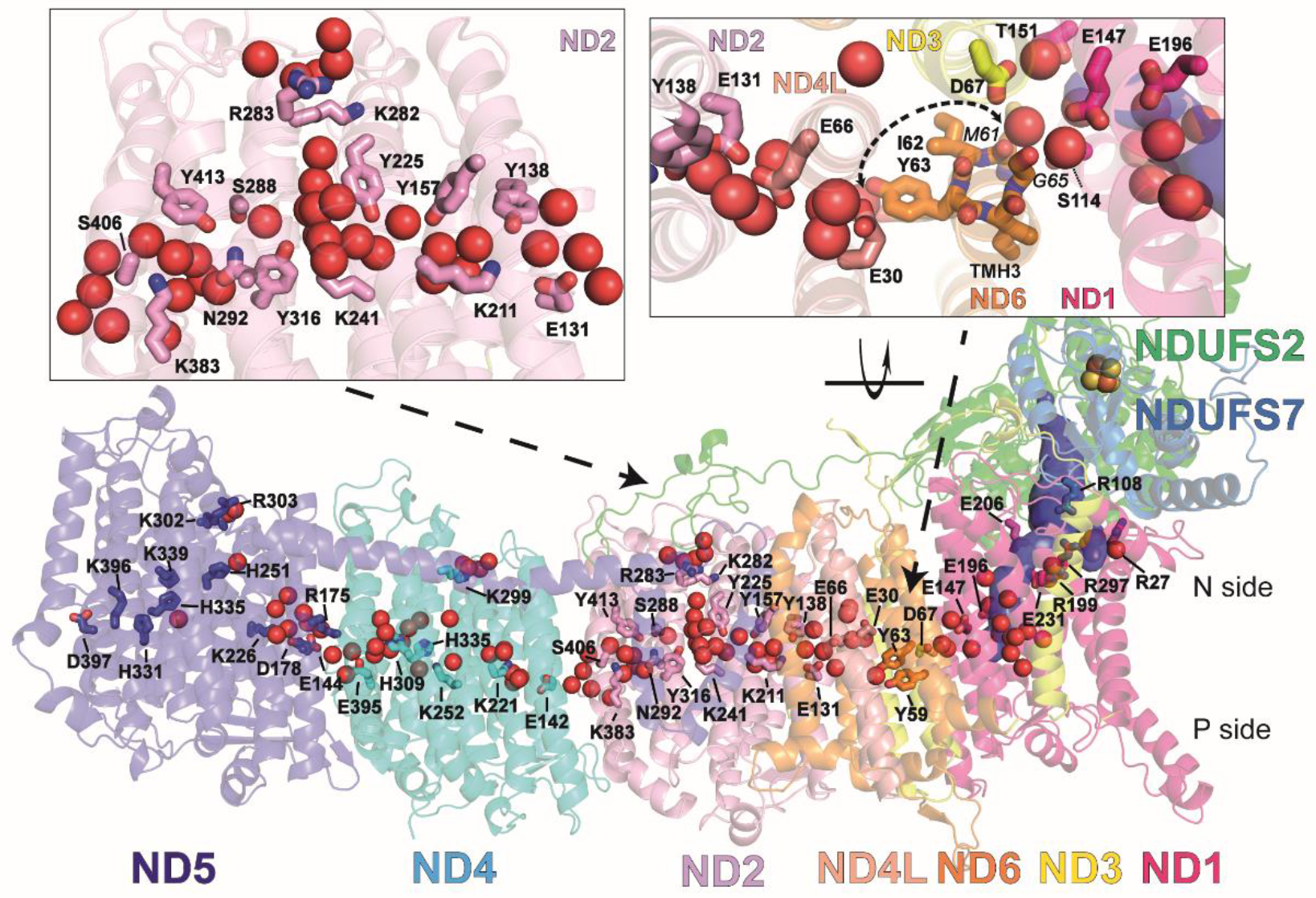
Protein-bound water molecules in the membrane arm. Side view of the central membrane and central subunits NDUFS2 and NDUFS7 of the Q module; the internal volume (blue surface) represents the E channel and Q access pathway; 99 water molecules (red spheres, see Figure S3) were resolved in the transmembrane region and at the entrance of putative proton uptake pathways at the N-side of the inner mitochondrial membrane. The left inset shows water molecules with interspersed hydrophilic and charged residues in ND2; the inset on the right shows a view of the π-bulge region of TMH3 of ND6 from the matrix side; residues interacting with water molecules with their carbonyl oxygens are labeled in italics. An ∼18 Å dry region (dashed line) separates the water cluster at ND1/ND3 from the water chain leading from ND4L to ND4. See also Figure S3-4 and S11.

Overall, a previous study on *Y. lipolytica* complex I (Figure S3) (Grba and Hirst, 2020) agrees with our structure, which is however substantially more complete and of significantly higher resolution, indicating the positions of 99 water molecules in the central membrane arm subunits (as compared to 34 in Grba and Hirst, 2020). Moreover, our high-resolution structure shows a clear proton input pathway from the N-side and a contiguous horizontal network of waters and connecting amino acid side chains in ND2. In the mammalian complex, subunit ND5 was more hydrated and similar numbers of water molecules were found in ND2 and ND4, but in ND2 a connection to the N side was not apparent (Kampjut and Sazanov, 2020).

### Atomistic molecular dynamics simulations of complex I

For a better insight into complex I dynamics, we performed atomistic MD simulations based on the 2.1 Å structure. We simulated two different protonation states of the enzyme (see Table S2, Figure S5 and computational methods). In state PN1, all titratable residues were modeled in their charged states, whereas in state PN2, the protonation state of sidechains was taken from the pK_a_ calculations (see Table S2 and computational methods). The MD simulations indicated that hydration of the E channel and the region between the channel and subunits ND3/ND6 depends on the charged state of the associated residues. Extensive hydration of this entire segment was observed only when all residues were modeled in their charged states (state PN1), whereas hydration dropped significantly in several regions upon charge neutralization (state PN2) of acidic residues (Figure S6A and S6B). A smaller number of water molecules observed in these map regions suggests a preponderance of titratable residues in their charge-neutral state, so that dynamic water molecules might not be sufficiently immobilized by interaction with the protein.

On the basis of pK_a_ estimates, we found that all four titratable residues of the central hydrophilic axis Glu131, Lys211, Lys241 and Lys383 in ND2 are charged, whereas in ND4, Glu395, Lys252 and Glu142 are predicted to be neutral. As expected, the higher overall charge in subunit ND2 resulted in a higher hydration level, compared to subunit ND4 (Figure 3A). PN2 state-based water occupancies are in excellent agreement with water positions from the high-resolution map for both the central hydrophilic axis and the N-side pathway in ND2 (Figure 3A, Figure S7). We analyzed the H-bonding connectivity in ND2 and found a remarkable, near-continuous hydrogen bond network capable of long-range lateral proton transfer along this section of the central membrane arm (Figure 3E). We note that the highly conserved residue Phe343 of TMH 11 in ND4 interrupts the water chain between the hydrated central axis and the N-side Lys299 (Figure 3B). We therefore modeled the conformational state of TMH 11 and charged state of the ND4 subunit in the same way as in ND2, where a higher level of hydration was observed between the N-side and the central axis (Figure 3A). Upon remodeling, we observed a rapid hydration of the region (Figure 3D) as in the ND2 subunit, suggesting a potential gating role of Phe343, which is conserved in all three antiporter-like subunits (Figure S4). Also, in the ND5 subunit, which differs from ND2 and ND4 at the sequence level (Figure S4), the N-side connection emerged at the conserved KR motif and continued towards His251 via Ser247 and stable water molecules (Figure 3C).

**Figure 3.**
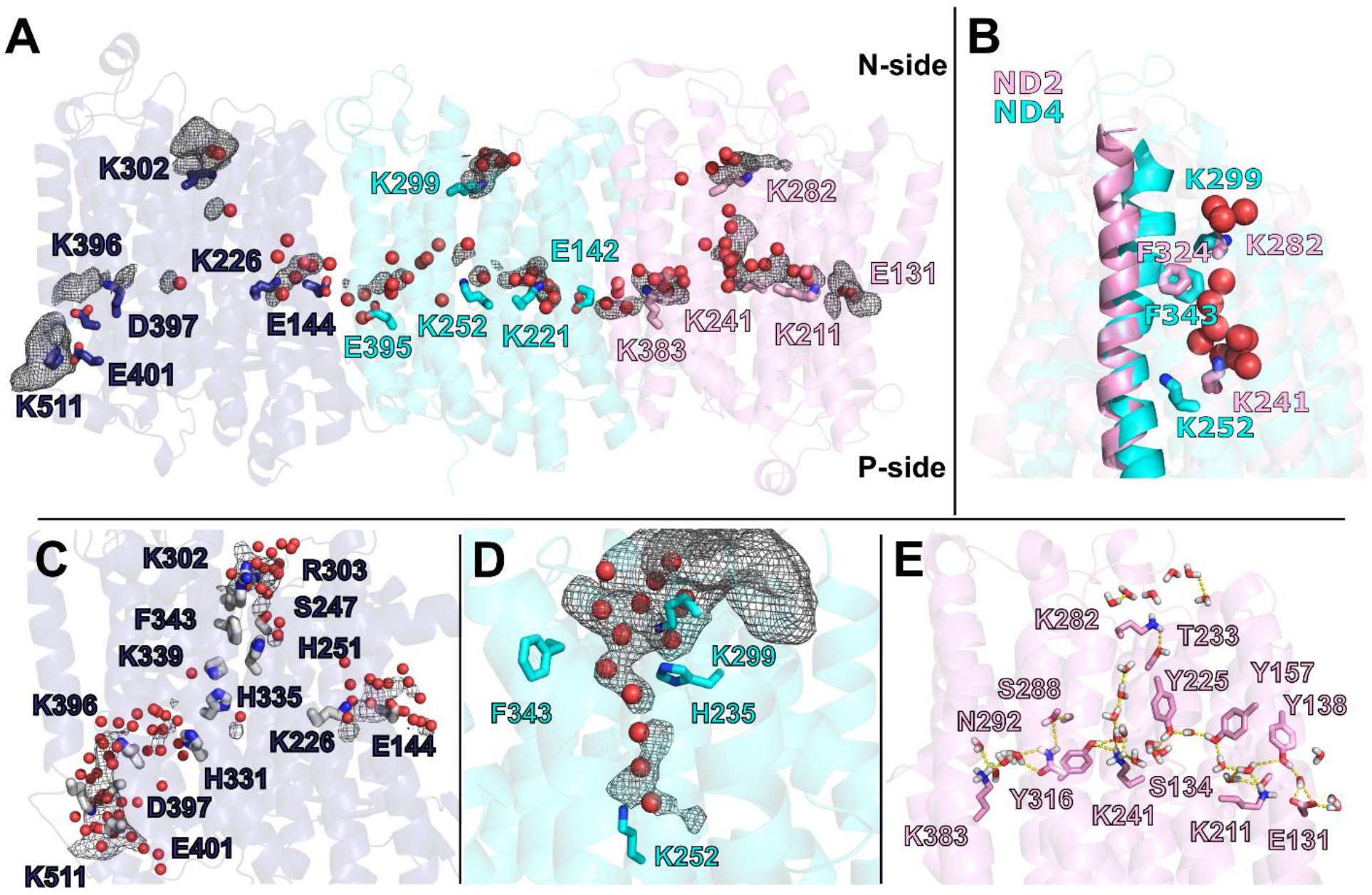
Hydration of antiporter-like subunits. (A) Water molecules resolved in the cryo-EM map (red spheres) and hydrated regions observed in MD simulations (grey mesh) in subunits ND5 (dark blue), ND4 (cyan) and ND2 (pink). The volumetric map (isovalue 0.15) is calculated from simulation PN2 by selecting water molecules within 6 Å from the labelled residues. (B) An overlay of ND2 and ND4 subunits with TMH11 and conserved Phe324/343, Lys282/299 and Lys241/252 residues highlighted. Due to the conformation of TMH11, Phe343 blocks the water-based N-side connection in the ND4 subunit. Resolved structural waters (red spheres) are from subunit ND2. (C) N- and P-side water/residue-based connections in the ND5 subunit from simulation PN1. The volumetric map (isovalue 0.2) is calculated by selecting MD-based water molecules (oxygen atoms, red spheres) within 6 Å of selected residues. (D) Water wire and volumetric map (isovalue 0.2) from the N-side to the central axis of the ND4 subunit after re-modeling of the conserved Phe343. (E) Simulation snapshot showing a continuous hydrogen bond network in subunit ND2 from simulation PN2. Yellow dashed lines indicate hydrogen bonds that meet the 3.6 Å distance and 30° angle criteria. Red and white sticks indicate water molecules. See also Figure S5-8.

However, the proton path was partly blocked between His251 and Lys339 by the conserved Phe343 residue, whereas from Lys339 onwards a continuous string of sidechains and water molecules formed towards the P-side. The conserved Phe in TMH 11 and its equivalent in the other two antiporter modules may thus play an important role in preventing a proton short-circuit between the two membrane surfaces. Overall, the MD simulations and structural data suggest strongly that three antiporter-like subunits differ from each other with respect to conformation, charge and hydration states.

Our cryo-EM structures and MD simulations revealed substantial hydration in the central segments of subunits ND2 and ND4. The hydrated paths in the middle of the subunits connect to the N-side of the membrane, as also observed in simulation studies of bacterial complex I (Di Luca et al., 2017; Haapanen and Sharma, 2017). A proton transfer path between the central axis and the P-side was not apparent in the structures or in any of the ND2/ND4 subunit simulations. In contrast, P-side connectivity was clearly observed in the ND5 subunit (Figure 3C). This is most likely due to the charged residues Asp397^ND5^, Glu401^ND5^, Lys511^ND5^ closer to the P-side that are only found in the ND5 subunit (Figure 3C and S4), but not in ND2/ND4. When the hydrophobic residues Phe396^ND4^ and Leu384^ND2^, which correspond to Asp397^ND5^ in the ND2/ND4 subunits, were changed to aspartate, we observed extensive hydration between the central segments of ND2/ND4 subunits and the P-side (Figure S6C-F). This suggests that hydrophobic residues in the ND2/ND4 subunits insulate the hydrated central axis from the P-side and prevent proton equilibration, thus conserving the electrochemical potential.

While there is only one conserved Lys396/Asp397 pair in the ND5 homologue of the bacterial enzyme, mitochondrial complexes contain an additional ion pair (Glu401 and Lys511), which further adds to the hydration of this region (Figure 3A, Figure 3C). The hydrated P-side path in ND5 encompasses a unique lipid-protein architecture in which accessory subunits NDUFB2 and NDUFB3 bend the lipid bilayer (Figure S8). This unique lipid-protein arrangement is confined to the ND5 subunit, and may thus be important in forming a proton-exit route that is characteristic of this subunit only. We suggest that perturbation of the lipid bilayer by the unique complex I architecture is crucial for proton pumping and exchange of the quinone substrate at subunits ND5 and ND1, respectively (see also (Haapanen et al., 2020; Parey et al., 2019).

### Complex I under turnover conditions

We previously reported an experimental strategy to capture complex I during steady-state NADH:decylbenzoquinone (DBQ) oxidoreductase activity for cryo-EM analysis (Parey et al., 2018). By enzymatic recycling in a minimal respiratory chain consisting of complex I and a quinol oxidase, this method ensures a sufficient supply of the oxidized hydrophobic substrate, and thus circumvents problems of low DBQ solubility at the concentrations required (>200 µM). We now improved the resolution from 4.5 Å to 3.4 Å, which enabled us to build a model of 7921 residues, all cofactors, 33 lipids and bound substrates in complex I under turnover conditions.

Overall, we observed a small shift of the P_D_ module towards the matrix side and towards the P_P_ module (Figure S9, Movie 2). The angle between peripheral arm and membrane arm decreased slightly by about ∼1 degree, corresponding to a 2-3 Å shorter distance between the distal ends of the two arms. At the junction of the two arms and towards the P_D_/P_P_ interface, the map density of the turnover map suggests a local loss of order and higher mobility of the protein structure in the interface region (Figure S9 E,F). Compared to mammalian complex I (Agip et al., 2018; Kampjut and Sazanov, 2020), the change in angle between peripheral arm and membrane arm in the *Y. lipolytica* complex I is moderate (Figure S9 C,D), most likely because the *Y. lipolytica* complex I is less flexible. Note that we recently observed larger rearrangements of the two arms comparable to mammalian complex I in an inactive mutant of *Y. lipolytica* complex I (Galemou Yoga et al., 2020b).

A clear new density in the NDUFV1 subunit enabled us to model an NADH molecule in electron-transfer distance to the flavin mononucleotide (Fig. S10A,B). Another clear density in the Q reduction site close to His95^FS2^ was modelled as the substrate quinone head group (Figure 4A, B). His95^FS2^ and Tyr144^FS2^ are thought to act as ligands for Q during electron transfer from FeS cluster N2 in NDUFS7 (Baradaran et al., 2013; Gutierrez-Fernandez et al., 2020; Sharma et al., 2015; Tocilescu et al., 2010). In our model the distance from the Q head group to Tyr144^FS2^ is 5-6 Å, which argues against a strong hydrogen bond. Met91^FS7^ points to the center of the Q head group, and Ser192^FS2^ is closer than 4 Å to one of the methoxy groups. The Q head group position agrees well with our previous observation (Parey et al., 2018) and with the recent structure of ovine complex I under turnover conditions (Fig. S10C) (Kampjut and Sazanov, 2020) but differs from the crystal structure of *T. thermophilus* complex I (Gutierrez-Fernandez et al., 2020) (Fig. S10D). A comparison with the empty Q binding site of complex I reveals structural changes in the β1β2 loop of NDUFS2 that harbors the functionally important residues His91^FS2^ and His95^FS2^ and at position 78 to 81 of NDUFS7 (Movie 3). This rearrangement narrows the distance between Gln90^FS2^ and Thr80^FS7^ and constricts the site.

**Figure 4:**
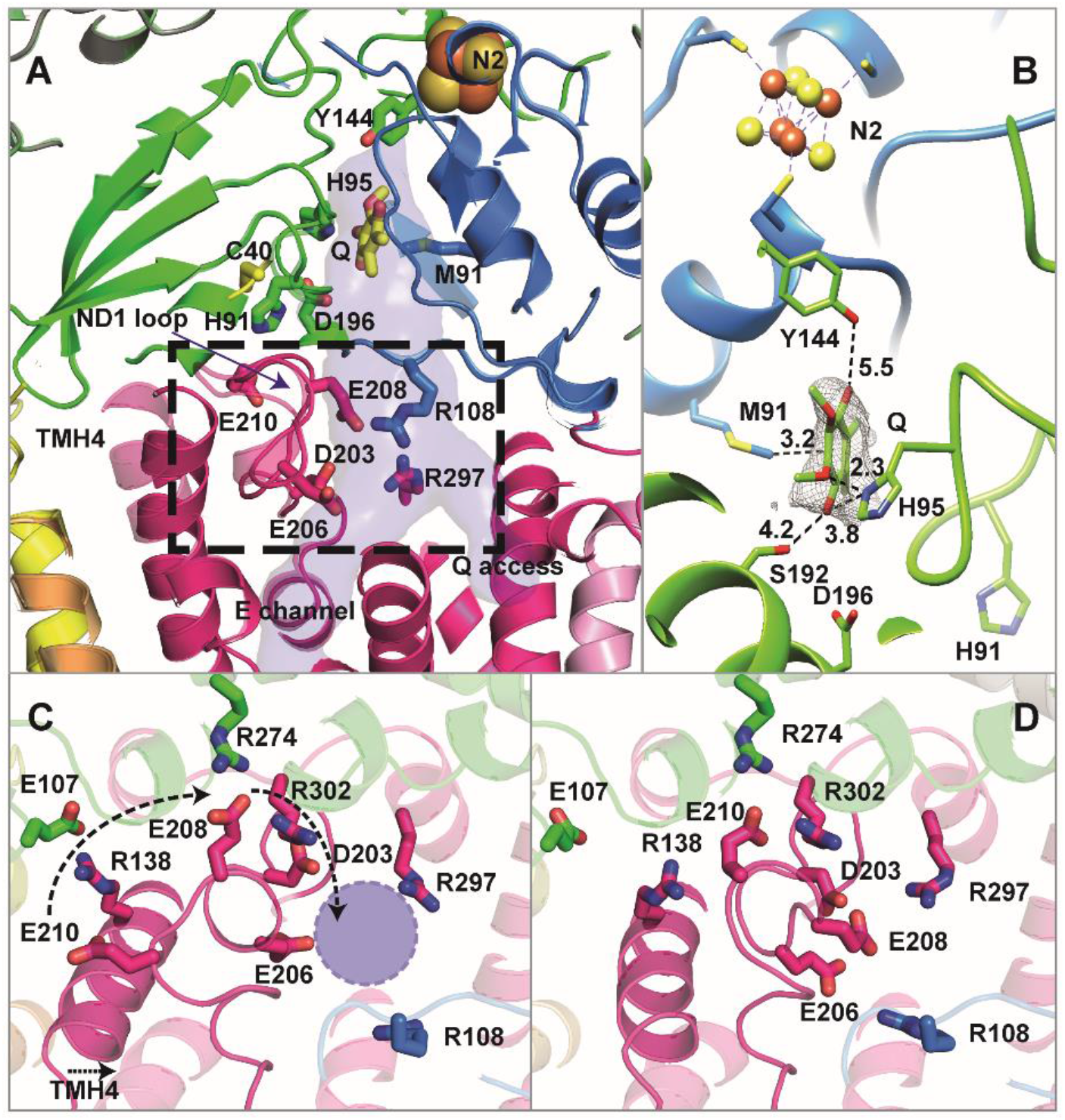
Q reduction site and conformational changes in ND1. (A) The Q reduction site near FeS cluster N2 is formed by subunits NDUFS2 (green) and NDUFS7 (blue) and is connected to the membrane by a tunnel crossing subunit ND1 (magenta); the branch point from where the E channel leads into the membrane arm interior is surrounded by protonatable residues. TMH4 and the TMH5-6 loop of ND1 (boxed area, see Figure 5) undergo a conformational change. (B) Binding of the Q head group in the Q reduction site; numbers indicate distances in Å. (C) Complex I in the D form, view from the matrix side towards TMH4 and the ND1 loop at the level of the branch point of E channel and Q access pathway (violet circle). (D) same view as in (C) for steady-state activity conditions. See also Figure S9-10 and Movie 2 and 3.

For mammalian complex I, different functional states involve the transition of TMH3 in ND6 from a π-bulge to a regular α-helix (Agip et al., 2018; Kampjut and Sazanov, 2020; Letts et al., 2019). It has recently been proposed that an uninterrupted water chain along the hydrophilic axis can form only when this helix is in the α-helical conformation (Kampjut and Sazanov, 2020). However, TMH3 of ND6 clearly forms a π-bulge helix in both our present structures of *Y. lipolytica* complex I. In our MD simulations based on the 2.1 Å resolution structure, we monitored transient changes in the region. Ile62^ND6^ was found to adopt a conformation that would tend to enhance local hydration (Figure S11). As a result, the dry ∼18 Å gap (Figure 2) is bridged by water molecules connecting the acidic residues of ND4L and ND3 (Figure S11B and S11C), so that large-scale rearrangements such as the π↔α transition are not required in *Y. lipolytica*. Instead, conformational changes at the sidechain level may be sufficient to achieve lateral proton transfer through this route.

### Conformational changes in membrane subunit ND1

Subunit ND1 forms the major part of the membrane/peripheral arm interface and therefore must play a key role in the coupling mechanism. We observed clear conformational changes in TMH4 and in the TMH5-6 loop of this subunit (Figure 4, Movie 3). The TMH5-6 loop contains the strictly conserved acidic residues Asp203^ND1^, Glu206^ND1^, Glu208^ND1^ and Glu210^ND1^ and is resolved in both our structures. In contrast, the loop is disordered in the open state (Kampjut and Sazanov, 2020; Letts et al., 2019) and in the deactive form (Agip et al., 2018) of mammalian complex I. Note that, while previous reports compared conformational changes of mouse and *Y. lipolytica* complex I (Grba and Hirst, 2020), we now describe the reorganisation of the ND1 TMH5-6 loop in one and the same enzyme complex.

In our high-resolution structure, the strictly conserved Arg274^FS2^ and Arg302^ND1^closely interact with Glu208^ND1^ (Figure 4C). Under turnover conditions, the C-terminal part of TMH4 bends, and the salt bridge between the strictly conserved Arg138^ND1^ at the helix kink and Glu107^FS2^ in subunit NDUFS2 opens (Figure 4D). Since this helix cannot bend without clashing with the ND1 loop near position 210, the loop must rearrange. The strictly conserved Glu210^ND1^ moves into the Arg274^FS2^/Arg302^ND1^ binding pocket, replacing Glu208^ND1^ which shifts more than 9 Å towards the Q tunnel. This results in a roughly triangular arrangement of Asp203^ND1^, Glu206^ND1^ and Glu208^ND1^ close to the branch point of Q tunnel and E channel. The conformation is similar to that observed in the closed state (Kampjut and Sazanov, 2020) and the active form (Agip et al., 2018) of mammalian complex I. The almost complete loss of activity after exchanging Glu107^FS2^ and Arg274^FS2^ by site-directed mutagenesis showed that both contact points between ND1 and NDUFS2 are critical for function (Fig. 5A). Note that ND1 cannot be mutagenized because it is encoded by mitochondrial DNA.

**Figure 5:**
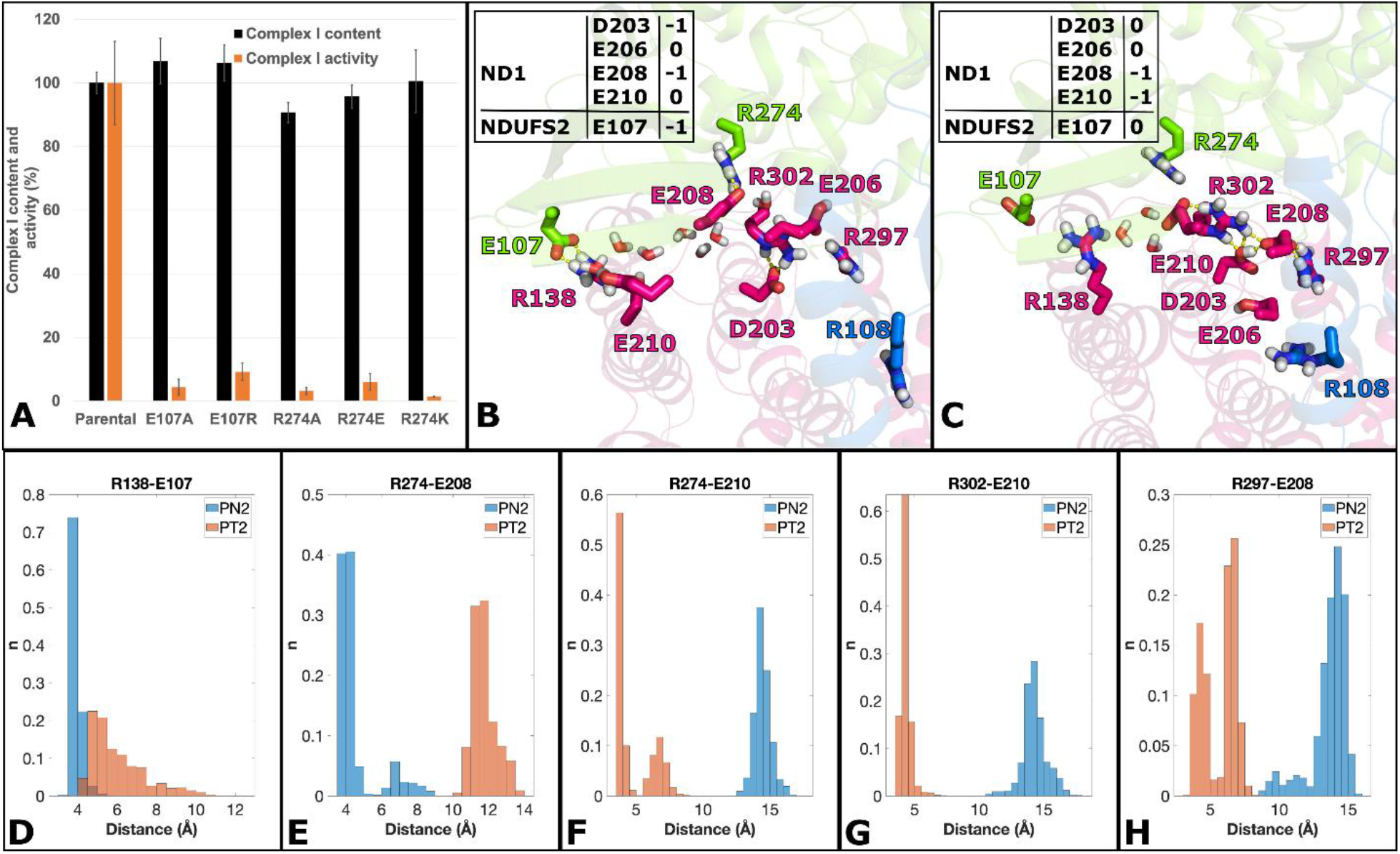
Mutagenesis and conformational rearrangement of key residues in subunits ND1 and NDUFS2. (A) Site-directed mutagenesis of subunit NDUFS2. Complex I assembly and inhibitor sensitive Q reductase activity of mutant complex I carrying exchanges at position Glu107 and Arg274 was tested in mitochondrial membranes (complex I assembly, black bars, 100% corresponds to 1.12 ± 0.04 μmol·min^−1^·mg^−1^ NADH: hexaaminruthenium oxidoreductase activity; activity, orange bars, 100% activity corresponds to 0.54 ± 0.07 μmol·min^−1^·mg^−1^ dNADH:decylubiquinone oxidoreductase activity). Simulation snapshots (B and C) of key interactions of selected residues from ND1 (pink), NDUFS2 (green) and NDUFS7 (blue) subunits from MD runs PN2 (B) and PT2 (C). The charged states of selected residues (based on pK_a_ calculation) are indicated in panels B and C. Stable interactions between residues differ between protonation states and are in agreement with the high-resolution structure (compare Figure 4, boxed area in panel A, note that viewing directions are different). Selected water molecules are shown as red-and-white sticks connecting the surface-exposed region (Arg138/Glu107/Glu210) to buried Asp203, next to the E channel. (D-H) Histograms of stable charged-charged interactions. Distances were measured from arginine CZ atoms to glutamic acid CD atoms throughout all trajectory data. Data are shown for both PN2 (blue) and PT2 (orange) MD runs. The bin size was 0.5 Å, and histograms are normalized such that the sum of the bar heights is equal to 1. See also Table S2 and S3.

To investigate the observed structural changes in the TMH5-6 loop, we performed MD simulations of the turnover structure (Table S2). In the first setup, PT1, all titratable amino acid residues were kept in the standard protonation state. In the second setup, PT2, protonation states were taken from pK_a_ calculations (Table S2). With these protonation states, the simulation recapitulated the interactions observed in the turnover structure very well (Figure 5). Hydration analysis revealed that the conserved acidic residues Glu107^FS2^/Glu210^ND1^ at the protein surface are connected by water molecules to the buried Asp203^ND1^ near the E channel, suggesting a novel proton uptake route (Figure 5). A comparison of calculated protonation states for the two conformations of the ND1 loop (Figure 5B,C) shows two remarkable differences. On the one hand, relocation and deprotonation of Glu210^ND1^ connects this residue to the positively charged binding pocket formed by Arg274^FS2^ and Arg302^ND1^. On the other hand, rearrangement of the loop and approach of the charged Glu208^ND1^ results in the uptake of a proton at Asp203^ND1^ at the entrance of the E channel (Figures 4, 5). In effect, a proton is transferred from the protein surface to the starting point of the route into the membrane arm through the novel pathway identified here.

It is interesting to note that the conformational changes in the ND1 loop are not isolated from the rest of the complex I structure. Movie 3 suggests that the rearrangements in ND1 are coordinated with changes in the Q reduction site and surrounding subunits, in agreement with earlier proposals (Galemou Yoga et al., 2019; Zickermann et al., 2015). Intriguingly, the bending motion of TMH4 coincides with movement of the P_D_ module (Movie 2). The mechanistic implications of this small movement are difficult to judge at this stage.

## Discussion

Most current mechanistic models of complex I propose that each of the antiporter-like subunits ND2, ND4 and ND5 encompasses a complete proton translocation pathway that connects the mitochondrial matrix (N-side) to the intermembrane space (P-side). This view was recently challenged (Kampjut and Sazanov, 2020). A fourth, as yet undefined proton pathway is thought to be either located at the ND3/4L/6 interfaces or to involve the E channel and a P-side connection in the P_P_ module. Our cryo-EM structures and MD simulations provide strong support for N-side connections in all three antiporter-like subunits, with a portal at the N-terminal end of TMH10. Our results also suggest a gating function of the conserved Phe in TMH11, which may work in conjunction with the recently proposed leucine gate in the loop region of discontinuous TMH7 (Grba and Hirst, 2020). In agreement with a study of the ovine complex (Kampjut and Sazanov, 2020), we find a P-side connection exclusively in ND5, which we show to comprise extra charged residues that increase hydration, and a unique lipid-protein interface. The number of charged residues in this region raises the possibility that some of them may trap and store protons, before they are ejected to the P-side. Note that the absence of other P-side connections in the membrane arm appears to be at variance with the residual proton translocation activity of a complex I mutant lacking the distal part of the membrane arm (Dröse et al., 2011). However, the structure of that mutant might be compromised by a leak pathway that does not exist in the wildtype. To account for a single P-side connection in the membrane arm, the present models for redox-linked proton translocation need to be revised.

The fact that complex I activity is fully reversible suggests that energy release at the Q site requires more than a single conformational change or charge rearrangement event. If ND5 is the only subunit through which protons are released to the P-side, it seems unlikely that the energy required for pumping of 4 H^+^ is transferred over a 200 Å distance by a single one-stroke mechanism. Instead, a mechanism with two separate electron/proton transfer events to Q via a semiquinone intermediate may be a better description. We propose that the conformational changes in ND1 that we describe here in detail are critical for injection of protons into the E channel and that these changes are linked with Q binding in the Q tunnel and its redox chemistry in the catalytic site (Figure 6). Protons injected into the E channel must travel the complete length of the hydrophilic axis, which would agree with our observation of a long contiguous connection towards ND4. It has recently been proposed that the π-to-α transition of TMH3 in ND6 is necessary for bridging a gap in the hydrophilic axis to enable proton movements between ND4L and ND1 (Kampjut and Sazanov, 2020) and that the TMH3 transition is tightly linked with a bending of TMH4 in ND1 (Grba and Hirst, 2020). Our data do not support either of these proposals because (i) structural changes in ND1, including a bending of TMH4, occur while the π-bulge is present and (ii) simulations show that a water/protein connection across ND6 can form after moderate changes in TMH3 at the side chain level. However, a transient formation of the α helix cannot be excluded at this stage.

**Figure 6:**
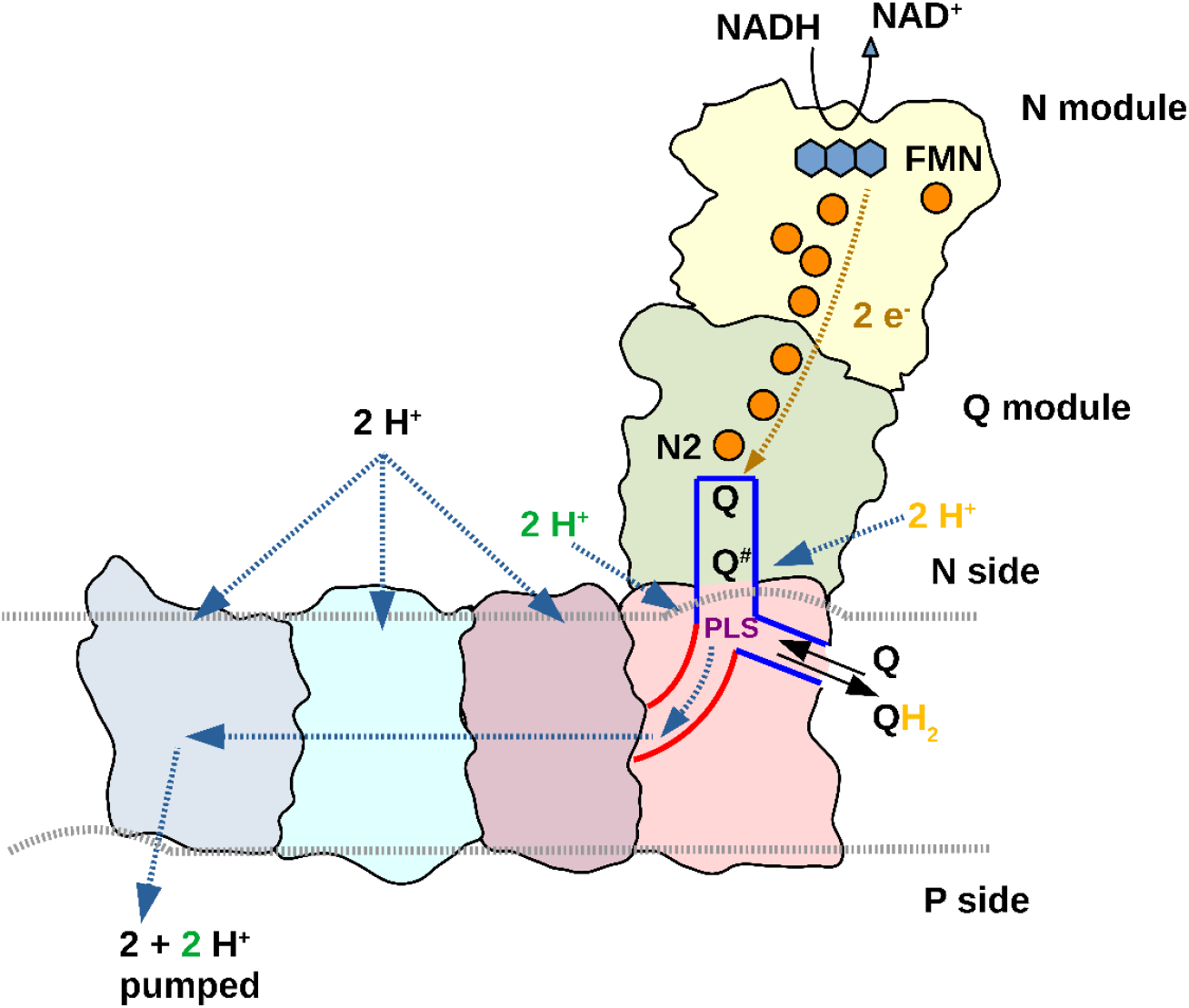
Model for redox-linked proton translocation by complex I. For electron transfer, Q binds in a site close to FeS cluster N2 and a negatively charged intermediate (Q^#^) is formed. Formation and stabilization of Q^#^ is associated with the loading of pumped protons (green) to a proton-loading site (PLS) by rearrangement of the ND1 loop (Fig. 4 and 5). The proton-loading site is situated at the junction of E channel (red) and Q tunnel (blue) in the vicinity of a second Q binding site identified previously (Haapanen et al., 2019; Parey et al., 2019). Translocation of Q^#^ to this second site might balance the positive charge of the proton stored in the proton-loading site. Uptake of a substrate proton (yellow) neutralizes the charge of Q^#^ and the resulting change in pK_a_ of the proton-loading site releases the proton into the E channel. The neutral Q intermediate moves back to the electron transfer site and the cycle repeats. Each PLS proton injected into the E channel is driven towards the antiporter-like subunits by a subsequently injected proton in the next cycle. The antiporter-like subunit can take up protons from the N-side. These protons are inserted into the hydrophilic axis and driven towards the only P-side exit at ND5 by repeated injection of protons from the E channel. Gating at the N-side by the conserved Phe in TMH11 (not shown) ensures directionality of proton movements and excludes short circuits. The experimentally determined pump stoichiometry of complex I is 4H^+^/2e^−^ (Galkin et al., 2006; Galkin et al., 1999; Jones et al., 2017; Wikström, 1984). In our model, two pumped protons are taken up from the E channel for each reaction sequence from Q to QH_2_. Therefore, the other two protons must be taken up via antiporter-like subunits. The odd parity between proton uptake and the number of proton-translocating subunits does not imply that only two of the antiporter-like subunits are active, but may indicate that complex I cycles through different functional states in which alternating N-side connections are silenced.

Based on these considerations, we propose a molecular mechanism of proton pumping in complex I, which resembles the well-documented proton pumping mechanism of cytochrome *c* oxidase, the terminal enzyme of the respiratory chain (Rich, 2017; Wikström et al., 2018). In cytochrome *c* oxidase, for every electron transfer to the oxygen reduction site, a proton is transferred to the proton-loading site, followed by substrate proton transfer to the active site, which then electrostatically drives the ejection of the *pumped* proton from the proton-loading site to the P-side of the membrane.

We propose that rearrangement of the ND1 loop causes the transfer of a proton from the N-side protein surface to the acidic cluster at the entrance of the E channel (Asp203^FS2^/Glu208^ND1^/Glu206^ND1^), which is the proton loading site of complex I (Figure 6). Movie 3 shows that the structural changes in ND1 are connected with rearrangements in the Q reduction site and are thus likely associated with Q binding and Q reduction. We suggest that every electron transfer to Q drives the uptake of one proton via the newly identified pathway and that the charge state of the proton loading site is balanced by the negative charge of the anionic Q intermediate that is formed in the electron transfer reaction. The transfer of a substrate proton for Q redox chemistry is energetically favorable and neutralizes the charge of the anionic Q intermediate. This protonation must be tightly controlled and may either occur at the Q binding site near N2 or at one of the other sites found in the Q tunnel (Haapanen et al., 2019; Parey et al., 2019; Warnau et al., 2018). A putative pathway for the delivery of substrate protons involving Glu39 of ND3 and His91 of NDUFS2 was recently discovered (Galemou Yoga et al., 2020b). Proton transfer to the Q intermediate nullifies the charge balance in the proton loading site and triggers the proton release into the E channel. The nearby positive charges of Arg297^ND1^ and Arg108^FS7^ may support deprotonation. Consistent with this notion, a reversal of the charge in mutant Arg108E^FS7^ has a strong impact on proton pumping activity of complex I (Galemou Yoga et al., 2019).

The set of mechanistic events described above would repeat for the second electron transfer to Q, feeding a total of two protons into the E channel. The functional dissection of electron and proton transfer steps to Q is consistent with the “two-state stabilization change” mechanism proposed earlier (Brandt, 2011). The two protons injected into the E channel are eventually released to the P-side after traversing the entire hydrophilic axis. For a stoichiometry of 4H^+^/2e^−^, two more protons must be taken up via N-side connections, and fed into the hydrophilic axis by the three antiporter-like subunits. We here present evidence for corresponding proton translocation pathways based on high-resolution structural data and MD simulations. We propose that the protons are driven towards ND5 and eventually to the P-side by successive injection of further protons into the hydrophilic axis via the E channel. This mechanism implies a functional asymmetry between ND2, ND4 and ND5 at each stage of the catalytic cycle, corresponding to distinct states e.g. for proton access and occlusion of a trapped proton to prevent back-leak reactions. We do observe clear differences in conformational, hydration and charge state between the three antiporter subunits and we propose that this asymmetry favours proton pumping at high proton-motive force. Details of the mechanism proposed here would require further experiments and theoretical work, but the proposed mechanistic resemblance between complexes I and IV of the respiratory chain may indicate how machines in bioenergetics have evolved similar coupling principles on the basis of electrostatics and conformational dynamics.

## Acknowledgements

This work was supported by the Deutsche Forschungsgemeinschaft (grant ZI 552/4-2 to VZ). VS acknowledges research funding from the Sigrid Jusélius Foundation, Academy of Finland (294652), University of Helsinki and the Magnus Ehrnrooth Foundation. We are thankful to the Center for Scientific Computing (CSC), Finland for high-performance computing support, specially acknowledging the pilot grand challenge project *Complexity2* (*mahti* supercomputer). We also acknowledge PRACE for awarding us access to Marconi100 at CINECA, Italy. We acknowledge excellent technical assistance by Karin Siegmund and Simone Prinz.

## Author contributions

KP purified and characterized complex I, prepared cryo-EM grids, acquired and processed cryo-EM data, analyzed data, drew figures and wrote the manuscript. JL performed all modeling and simulations, analyzed data, drew figures and wrote the manuscript. DJM collected cryo-EM data. AD analyzed data and drew figures. OH analyzed data and drew figures. EGY performed mutagenesis and analyzed mutants. HX provided quinol oxidase. WK provided cryo-EM infrastructure, supervised the cryo-EM work and wrote the manuscript. JV analyzed data, built the models, drew figures and wrote the manuscript. VS analyzed data and interpreted its mechanistic implications, supervised modeling and simulation work, drew figures and wrote the manuscript. VZ designed the study, interpreted the mechanistic implications of the structures, drew figures, and wrote the manuscript.

## Declaration of interest

The authors declare no competing interests.

## Experimental methods

### *Site-directed mutagenesis of complex I subunits from* Y. lipolytica

All point mutations were generated by inverse PCR mutagenesis of PUB26/NUCM plasmid for transformation in *E. coli*. The desired point mutation was introduced using appropriate primer pairs. For site directed mutagenesis, 5’-hydroxyl ends of primers were phosphorylated using T4 polynucleotide kinase prior to PCR. The PCR reaction mixture contained the template DNA (10 ng of plasmid DNA harboring the NUCM gene), both primers (forward primer and reverse primer; 0.5 μM each), dNTPs (200 μM), 1x Phusion HF buffer and Phusion DNA polymerase (2 U) in a final volume of 50 μl. The remaining un-mutated template DNA was removed by digestion with DpnI overnight at 37°C. Digested DNA fragments were separated onto a 1% agarose gel stained with peqGREEN from Peqlab at 120V for 1h. Linear DNA (∼11 kb) were cut out of the agarose gel after electrophoresis and isolated using the NucleoSpin Gel and PCR Clean-up kit from Macherey & Nagel according to the manufacturers’ instructions. Linear PCR products were ligated by blunt-end ligation using the Fast-link DNA ligation kits from Epicentre. 5μl of the ligation reaction were used for the transformation of chemically-competent *E. coli* XL-Gold cells. 1 ml LB was added to the mixture and incubated for 1h at 37°C on a rotating wheel.

Finally, 50-100 μl was spread on ampicillin-containing LB plates. The plates were incubated overnight at 37°C. On the following day, 5 clones were picked for each mutant and cultivated in 5 ml of LB medium containing ampicillin for 16h at 37°C. After overnight culture, plasmid DNA was isolated from *E. coli* cells using the NucleoSpin Plasmid kit from Macherey & Nagel and according to manufacturers’ instructions. After sequencing, plasmids DNA containing the desired mutation were transformed into *Y. lipolytica* strain Δ*nucm* using the lithium acetate method. Briefly, cells from 0.5–1 ml of an overnight culture were spun down, washed with 0.9% NaCl and centrifuged again. The cells were re-suspended in 90 µl of 50% PEG4000 with 100 mM lithium acetate pH 6, 100 mM DTT and 0.25 mg/ml ssDNA. 200–300 ng of plasmid DNA were added for each transformation mixture, vortexed and incubated at 39 °C for 60 min. After transformation, the mixture was spread onto a selective YPD plate containing 50 µg ml^−1^ hygromycin and incubated for 3 to 4 days at 28 °C.

### Preparation of mitochondrial membranes

Preparation of mitochondrial membranes of parental and mutant strains was carried out on a small scale as described (Galemou Yoga et al., 2019). Pellets of 5-8 g of cells (wet weight) were resuspended in 10 ml mito-buffer (600 mM sucrose, 20 mM MOPS, 1 mM EDTA, pH 7.2) supplemented with 2 mM PMSF and broken by vortexing with 10 g glass beads (0.25–0.5 mm) in a series of 15 × 1 min with 1 min resting intervals on ice. Cells debris and glass beads were removed by centrifugation at 3238 × *g* for 30 min at 4 °C. Mitochondrial membranes were sedimented from supernatant by ultracentrifugation at 147,642 × *g* for 1 h at 4 °C. Finally, the pellets were resuspended in 0.5–1 ml mito-buffer with 5 mM PMSF and homogenized. Aliquots of the membrane suspension were immediately frozen in liquid nitrogen and stored at −80 °C.

### Complex content and activity in mitochondrial membranes

Complex I content in mitochondrial membranes was determined as NADH:HAR oxidoreductase activity. This unphysiological activity reflects the amount of fully assembled complex I in a preparation and was determined spectrophotometrically at 340 nm wavelength (ε = 6.22 mM^−1^ cm^−1^) by measuring the initial oxidation rate of NADH. The reaction was started by the addition of 20-40 µg protein of mitochondrial membranes to the HAR buffer (250 mM sucrose, 20 mM HEPES, 0.2 mM EDTA, 2 mM KCN; pH 8) containing 200 µM NADH and 2 mM HAR.

Complex I activity in mitochondrial membranes was determined spectrophotometrically at 340 nm wavelength (ε = 6.22 mM^−1^ cm^−1^) by measuring the physiological electron transfer activity of deamino-NADH (dNADH) to the short chain ubiquinone analogue DBQ. dNADH is specific to complex I and is used in the complex I activity assay to exclude oxidation by alternative dehydrogenases. The reaction was started by the addition of 100 µM dNADH to a solution of DBQ buffer (20 mM MOPS-Na; 50 mM NaCl; 2 mM KCN; pH 7.2) containing 60 µM DBQ and 20–40 µg protein of mitochondrial membranes. The reaction was stopped by the addition of 2 µM complex I inhibitor 2-decyl-4-quinazolinylamine (DQA). The inhibitor-insensitive activity was subtracted from the initial measurement and the result was normalized to complex I content to allow comparison between different preparations. All activity measurements were performed at 30 °C in a Shimadzu UV-2450 spectrophotometer. For each mutant two biological replicates of mitochondrial membranes were analyzed, and each measurement was carried out in duplicate.

### Complex I purification and characterization

Complex I was purified in the detergent lauryl maltose neopentyl glycol (LMNG) by His-tag affinity and size exclusion chromatography as described (Parey et al., 2019), with an additional polishing step on a Mono-Q ion exchange column. Inhibitor sensitive NADH:decylubiquinone oxidoreductase activity of purified complex I was measured as described (Parey et al., 2019) but with 0.0015 % LMNG in the assay buffer.

### In vitro *assay of a minimal respiratory chain*

The *in vitro* assay of a minimal respiratory chain was carried out in 2-ml glass vials with stirring in a water bath at 10 °C. The reaction vial was filled with 20 mM Tris-HCl (pH 7.2), 100 mM NaCl and 0.025% LMNG, followed by the addition of 200 μM 2,6-dimethoxy-1,4-benzoquinone (DBQ), 2 μM complex I and 1 μM cytochrome *bo*_3_ ubiquinol oxidase (complex IV) to a final volume of 600 μl. The reaction was started by adding 2 mM reduced nicotinamide adenine dinucleotide (NADH) and inhibited by the addition of 2-decyl-4-quinazolinyl amine (DQA) or potassium cyanide (KCN). Oxygen consumption was determined polarographically using a Clark-type oxygen electrode (OX-MR; Unisense) connected to a picoammeter (PA2000 Multimeter; Unisense). Signals were converted with an A/D converter (ADC-216; Unisense) and then recorded with the software Sensor Trace Basic 2.1 supplied by the manufacturer.

### Cryo-EM structures

Complex I samples were applied at a concentration of 1.5 mg/ml in 20 mM Tris/HCl, pH 7.2, 100 mM NaCl, 1 mM EDTA and 0.025% LMNG to freshly glow-discharged 1.2/1.3 holey carbon grids (Protochips, USA). Grids were blotted for 12–14 s in a Vitrobot Mark IV (Thermo Fisher Scientific Inc., USA) at 10 °C and 70% humidity (drain and wait time 0 s, blot force -2), plunge-frozen in liquid ethane and stored in liquid nitrogen until further use. Cryo-EM data were collected automatically using EPU software (Thermo Fisher Scientific Inc., USA) on a FEI Titan Krios microscope (Thermo Fisher Scientific Inc., USA) at 300 kV equipped with K3 Summit detector (Gatan, USA) operating in counting mode. Cryo-EM images were acquired at a nominal magnification of 160,000x with a calibrated pixel size of 0.5316 Å, at a defocus range from −0.8 to −2.2 μm, with an exposure time of 3 s, and a total electron exposure on the specimen of ∼50 e^−^/Å^2^.

A set of 21,770 dose-fractionated micrographs from four independently collected data sets were subjected to motion correction and dose-weighting by MotionCor2 (Zheng et al., 2017). The micrograph-based contrast transfer function (CTF) was determined by Gctf (Zhang, 2016). CTF-corrected images were used for further analysis with the software package RELION3.1 (Zivanov et al., 2018). Particles were picked using Autopick within RELION3.1. A total of 1,078,966 particles was extracted with a box size of 600×600 pixels. Extracted particles from each data set were subjected to reference-free 2D classification with four-fold binned data to remove false positives and imperfect particles. A further 3D classification step with a previous cryo-EM map of complex I (EMD-4873) (Parey et al., 2019) as an initial reference was applied. The 3D classes from each data set were combined and used for auto-refinement, CTF refinement and Bayesian polishing in RELION3.1. (Zivanov et al., 2018). The final map from 178,960 particles at 2.4 Å resolution was sharpened using an isotropic B-factor of –47 Å^2^ (Fig. S1, Table 1) and the local resolution was estimated with ResMap (http://resmap.sourceforge.net) (Kucukelbir et al., 2014) (Fig. S1). All resolutions were estimated using the 0.143 cut-off criterion (Rosenthal and Henderson, 2003) with gold-standard Fourier shell correlation (FSC) of two independently refined half maps (Scheres, 2012). Density modification with *phenix.resolve_cryo_em* (Terwilliger et al., 2020) was carried out using two half-maps together with the FSC-based resolution and the post-processed map. This resulted in a significant improvement of the map quality, at a final resolution of 2.12 Å.

The cryo-EM structure of *Y. lipolytica* complex I (PDB id: 6rfr) was used as template and fitted to the map with COOT (Emsley et al., 2010). The structure was manually rebuilt in COOT and refined using *phenix.real_space_refine* (Afonine et al., 2018). Water molecules were built automatically with *phenix.douse* and checked manually. The “check/delete water” function in COOT was used to verify the molecules with clear map features and appropriate hydrogen bonding and geometry. Refinement and validation statistics are summarized in Table S1.

For cryo-EM studies of complex I under turnover, 4 μM of complex I incubated with 400 μM DBQ was mixed 1:1 with 2 μM *bo*_3_-type ubiquinol oxidase and 4 mM NADH. 3 μl of the mixture was applied to freshly glow-discharged 1.2/1.3 holey carbon grids (Protochips, USA) as described above. Cryo-EM data were collected automatically at 300 kV using EPU (Thermo Fisher Scientific Inc., USA) on an FEI Titan Krios microscope (Thermo Fisher Scientific Inc., USA) equipped with a Gatan K2 in movie mode at a calibrated magnification of 1.077 Å pixel^−1^ (105,000x). Cryo-EM images were acquired at a total exposure of 40 e^−^/ Å^2^ at −0.8 to −3.2 μm defocus.

CTF parameters were estimated by Gctf (Zhang, 2016) within the RELION3.0 workflow (Zivanov et al., 2018). A total of 124,092 particles were extracted from 4776 motion-corrected micrographs. The particles were subjected to reference-free 2D classification and sorted by 3D classification, which resulted in a final dataset of 54,863 particles that yielded a 3.4 Å map after CTF refinement and Bayesian polishing. An isotropic B-factor of –62 Å^2^ was applied and the local resolution was estimated with ResMap (Fig. S1).

The cryo-EM structure of complex I in the D form (Parey et al., 2019) was used as template and the structure was fitted into the map using *phenix.real_space_refine* in combination with rigid-body refinement. Critical loops were rebuild and the model was checked with COOT (Emsley et al., 2010).

Figures were made with Chimera (Pettersen et al., 2004) and PyMOL (Schrödinger and DeLano, 2020). Movies were made with ChimeraX (Goddard et al., 2018).

## Computational methods

Atomistic molecular dynamics simulations were performed on the complete 2.1 Å and 3.4 Å structures of *Y. lipolytica* complex I, including all resolved lipids, ligands, and water molecules. The protein was embedded into a membrane of 50% POPC, 35% POPE and 15% cardiolipin, generated with CHARMM-GUI (Jo et al., 2008). The system was solvated with TIP3 water and 100 mM of Na^+^/Cl^−^ ions. The force field for protein, lipids, water, and ions was CHARMM36 (Klauda et al., 2010; MacKerell Jr et al., 1998) and corresponding force field parameters were used for oxidized Q1 (Galassi and Arantes, 2015), FMN (Freddolino et al., 2006), FeS clusters (Chang and Kim, 2009), NADPH (Pavelites et al., 1997), and ZMP (Jo et al., 2008). One model system was constructed each for the two structures with all titratable residues either kept in their standard protonation states (MD runs PN1 and PT1) or with the protonation states of titratable residues fixed based on pK_a_ values estimated with Propka (Olsson et al., 2011) (MD runs PN2 and PT2; see Table S2). The total system size was around 1.3 million atoms (Figure S5).

Gromacs v2020.3 (Abraham et al., 2015) was used to perform the system minimization, equilibration and production MD runs. An energy minimization step was applied to release strain from the system, with restraints on protein heavy atoms and lipid phosphorus atoms, followed by a 100 ps NVT equilibration using velocity rescale algorithm (Bussi et al., 2007) with the same restraints. The final equilibration was a 10 ns NPT step using velocity rescale algorithm and Berendsen barostat (Berendsen et al., 1984), with restraints only on protein backbone atoms. In the production run, all restraints were released and the Nose-Hoover thermostat (Hoover, 1985; Nosé, 1984) and Parinello-Rahman barostat (Parrinello and Rahman, 1981) were applied at a temperature of 310 K and a pressure of 1 atmosphere. The LINCS algorithm was used to achieve a 2 fs timestep (Hess, 2008). Trajectory analysis was performed using Visual Molecular Dynamics (VMD) (Humphrey et al., 1996). The total simulation time was 4.8 μs.

## Data availability

Cryo-EM maps of D form and turnover structure have been deposited in the electron microscopy data bank (EMDB) under accession code EMD-12742 and EMD-12741, respectively. The atomic models have been deposited in the protein data bank (PDB) under accession codes 7o71 and 7o6y, respectively.

## Notes

### Competing Interest Statement

The authors have declared no competing interest.

