## Supplemental Material for "High-resolution structure and dynamics of mitochondrial complex I – insights into the proton pumping mechanism"

**Supplemental Tables**

**Table S1. Related to Figure 1. Data collection, refinement and model statistics.**

| | **D form** | **turnover** |
|---|---|---|
| **Data collection** | | |
| Microscope | FEI Titan Krios | FEI Titan Krios |
| Camera | Gatan K3 Summit | Gatan K2 Summit |
| Voltage (kV) | 300 | 300 |
| Nominal magnification | 165,000x | 105,000x |
| Calibrated pixel size (Å) | 0.5316 | 0.828 |
| Electron exposure ($e^-$/Å$^2$) | 50.0 | 40.0 |
| Exposure time total (s) | 3 | 8 |
| Number of frames per image | 50 | 40 |
| Defocus range (µm) | -0.8 – -2.2 | -0.8 – -3.2 |
| **Image processing** | | |
| Motion correction software | *MotionCor2* | *MotionCor2* |
| CTF estimation software | *Gctf* | *Gctf* |
| Particle selection software | *RELION3.1* | *RELION3.0* |
| Micrographs (no.) | 21,770 | 4,776 |
| Initial particle images (no.) | 1,078,960 | 124,092 |
| Final particle images (no.) | 178,960 | 54,863 |
| Applied *B*-factor (Å$^2$) | -47 | -62 |
| Final resolution Relion (Å) | 2.42 | 3.41 |
| Final resolution denmod (Å) | 2.12 | - |
| **Refinement statistics** | | |
| Initial model | 6rfr | 6rfr |
| Modeling software | *COOT, PHENIX* | *COOT, PHENIX* |
| Protein residues | 8035 | 7921 |
| Ligands | 55 | 56 |
| Waters | 1617 | - |
| Map CC (volume) | 0.8181 | 0.8158 |
| RMS deviations | 0.005 | 0.007 |
| Bond lengths (Å) | | |
| Bond angles (°) | 0.797 | 0.878 |
| Ramachandran plot | | |
| Outliers (%) | 0.05 | 0.09 |
| Favored (%) | 96.88 | 96.46 |
| Rotamer outliers (%) | 2.88 | 0.56 |
| Molprobity score | 2.06 | 1.91 |
| All-atom clashscore | 9.81 | 14.92 |
| **PDB ID** | **7o71** | **7o6y** |

**Table S2. Related to Figures 3 and 5. MD simulation setups.**

| | Structure for simulation setup | Charge state | Simulation time x no. of replicas |
|---|---|---|---|
| **PN1** | 2.1 Å | Standard | 400 ns x 3 |
| **PN2** | 2.1 Å | Propka-based | 400 ns x 3 |
| **PN3** | 2.1 Å L384D (ND2) F396D (ND4) D397A (ND5) | Standard | 400 ns x 1 |
| **PN4** | 2.1 Å F343 (ND4) modelled to F324 (ND2) position | Propka-based[1] | 400 ns x 1 |
| **PT1** | 3.4 Å | Standard | 400 ns x 1 |
| **PT2** | 3.4 Å | Propka-based | 400 ns x 3 |

[1]ND4 charges were set to replicate the ND2 charges calculated by Propka

**Table S3. Related to Figures 3 and 5. Amino acid residues with non-standard protonation states (neutral charge state of Asp, Glu and Lys and positive charge state of His) based on Propka calculation.**

| Subunit | 2.1 Å structure | 3.4 Å structure |
| --- | --- | --- |
| NDUFS1 | D259<br>D295<br>D377<br>D556<br>E355<br>E420 | D146<br>D295<br>D377<br>D556<br>E355<br>E420 |
| NDUFV1 | D120<br>D420<br>E121<br>E142<br>E210<br>E291<br>K276 | H378<br>D120<br>D420<br>E78<br>E121<br>E210<br>E291<br>K276 |
| NDUFS2 | D196<br>D331<br>D363<br>D465<br>E151<br>E376<br>E379<br>E463 | D196<br>D331<br>D465<br>E107<br>E151<br>E165<br>E208<br>E218<br>E379<br>E404<br>E463 |
| NDUFS3 | E179<br>K93 | E179<br>E181<br>E227<br>E237<br>K93 |
| NDUFV2 | H28<br>E211<br>K68 | H28<br>E211 |
| NDUFS8 | E186 | D163<br>E137<br>E186<br>E195 |
| ND1 | E147<br>E196<br>E206<br>E210<br>E231<br>K285 | D203<br>E101<br>E147<br>E196<br>E206<br>E231<br>K285 |
| ND2 | D39<br>D69<br>K282 | D39<br>D69<br>K282 |
| ND3 | D67<br>E39<br>E69 | D67<br>E39<br>E69 |

| | | |
|---|---|---|
| ND4 | E142<br>E395<br>K252<br>K299 | E58<br>E142<br>E237<br>E395<br>K252 |
| ND5 | D178<br>D397<br>E80<br>E503<br>K339<br>K581 | D501<br>E80<br>E144<br>E271<br>E401<br>E453<br>K339<br>K581 |
| ND6 | K182 | - |
| ND4L | D49<br>E30<br>E66 | E30<br>E66 |
| NDUFA9 | H76<br>D210<br>E231 | D210<br>E260 |
| NDUFA5 | H108<br>E92<br>E109<br>E110 | E92<br>E110 |
| NDUFS4 | E72<br>K111 | K111 |
| NDUFA12 | H78<br>D39 | H78<br>D39 |
| NDUFA6 | - | E82 |
| NDUFA2 | K41 | H62<br>E46 |
| NDUFA11 | - | H160 |
| NDUFA13 | H114<br>D7<br>E77 | D7<br>E77 |
| NDUFS5 | E53 | E18<br>E53 |
| NDUFA1 | H29 | H29<br>E51<br>E83 |
| NDUFB11 | - | E189 |
| NDUFB8 | D74<br>E33 | D59<br>D74<br>E33 |
| NDUFB7 | E58 | H49<br>E55<br>E58 |
| NDUFAB1 | - | E94<br>E100 |
| NDUFB10 | E39 | - |

**Supplemental Figures**

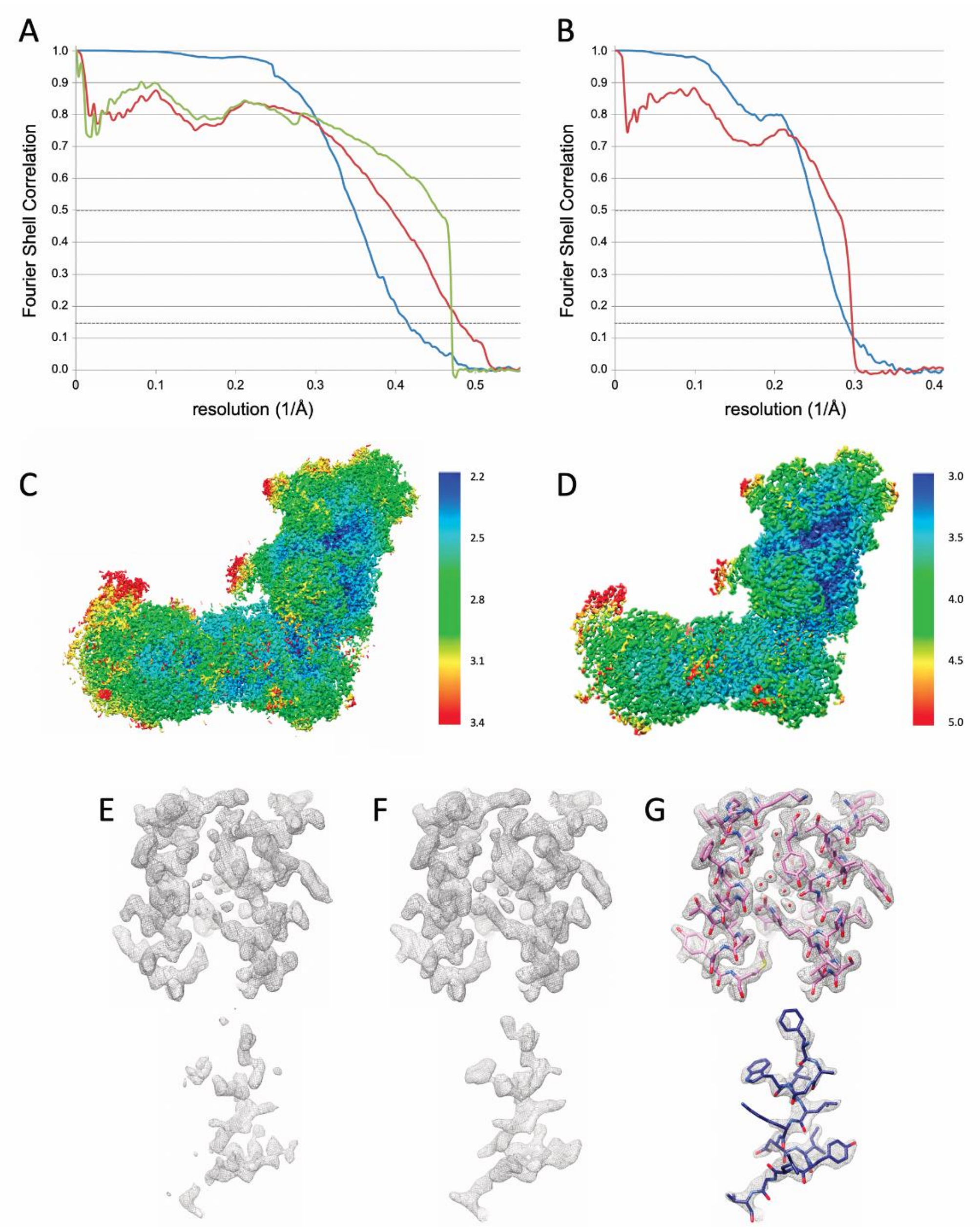


**Figure S1. Map validation. (A)** FSC plots for resolution estimation and model validation of complex I in the D form. Gold-standard FSC plot between two half-maps separately refined in RELION3.1 (blue) (Zivanov et al., 2018) indicates a resolution of 2.4 Å (0.143 threshold). Map-to-model FSC for the final refined model and the RELION map (red) and the density modified map (green) indicate a resolution of 2.5 and 2.2 Å, respectively (0.5 FSC criterion). **(B)** FSC plots for complex I under turnover. Gold-standard FSC (blue curve): 3.4 Å, map-to-model FSC (red): 3.6 Å. **(C)** D form and **(D)** turnover cryo-EM map colored by local resolution as determined in RELION. Color scale in Å. **(E-G)** Details of the wt cryo-EM density before **(E)** and after **(F)** density modification in Phenix (Terwilliger 2020). **(G)** density modified map with fitted model. Red spheres: water molecules. The procedure improved the average resolution of the map from 2.4 to 2.1 Å. It had subtle effects in high-resolution areas, especially improving water densities (top), while improving the continuity of less well resolved features (bottom).

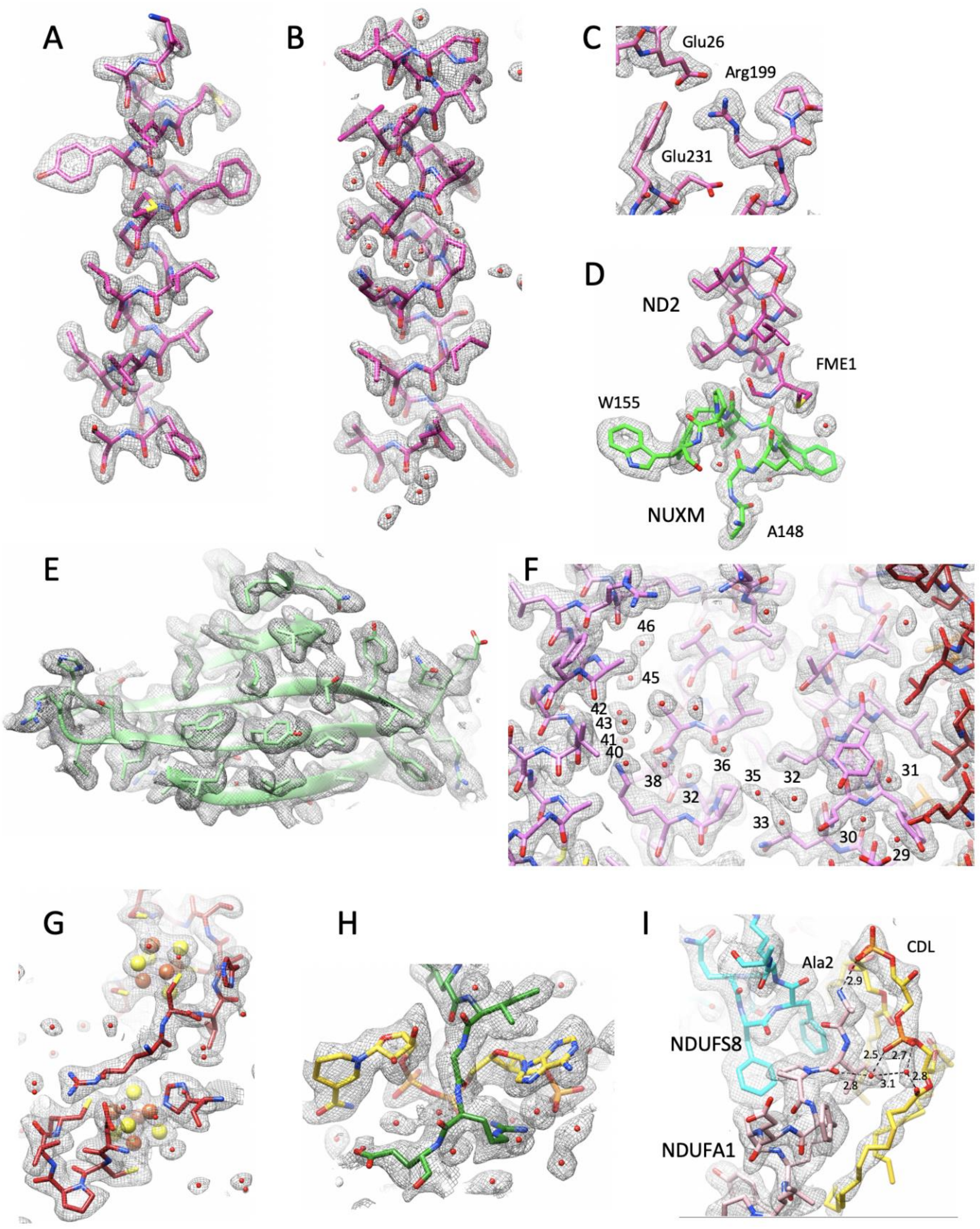


**Figure S2. Examples of cryo-EM density of complex I at 2.1 Å resolution. (A)** TMH ND2 152-171. **(B)** TMH ND2 229-249 with π-bulge. **(C)** salt bridge in ND1 with complete density for Glu26. **(D)** N-terminus of ND2 with formylmethionine, interacting with NUXM. **(E)** β sheet in NDUFS3. **(F)** Water molecules in the hydrophilic axis, labelled as in Figure S3. **(G)** FeS clusters in NDUFS1. **(H)** NADPH in NDUFA9. **(I)** Cardiolipin, coordinated by N-terminal Ala of NDUFA1; distances in Å are indicated.

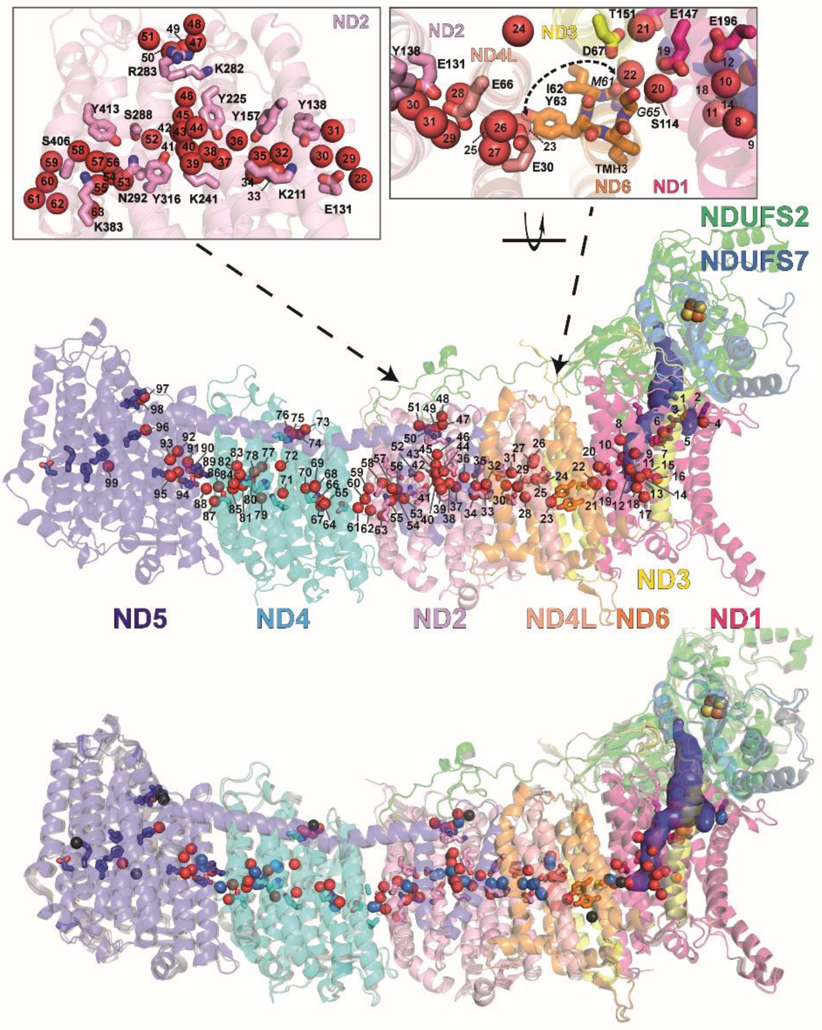


**Figure S3. Water molecules in the membrane arm of *Y. lipolytica* complex I.** Side view of central membrane bound subunits and central subunits NDUFS2 and NDUFS7 of the Q module as shown in Figure 2 with numbering scheme for the 99 water molecules (red spheres) resolved in the transmembrane region and at the entrance of putative proton uptake pathways. The upper left and upper right panel show detailed views of ND2 and centered around TMH3 of ND6, respectively. In the lower panel an overlay with the 2.7 Å structure of (Grba and Hirst, 2020)(PDB ID: 6yj4; cartoon, gray) with water molecules in central membrane arm subunits (waters exclusively found in 6yj4, black spheres; waters exclusively found in this work, red; overlapping positions, blue) is shown.

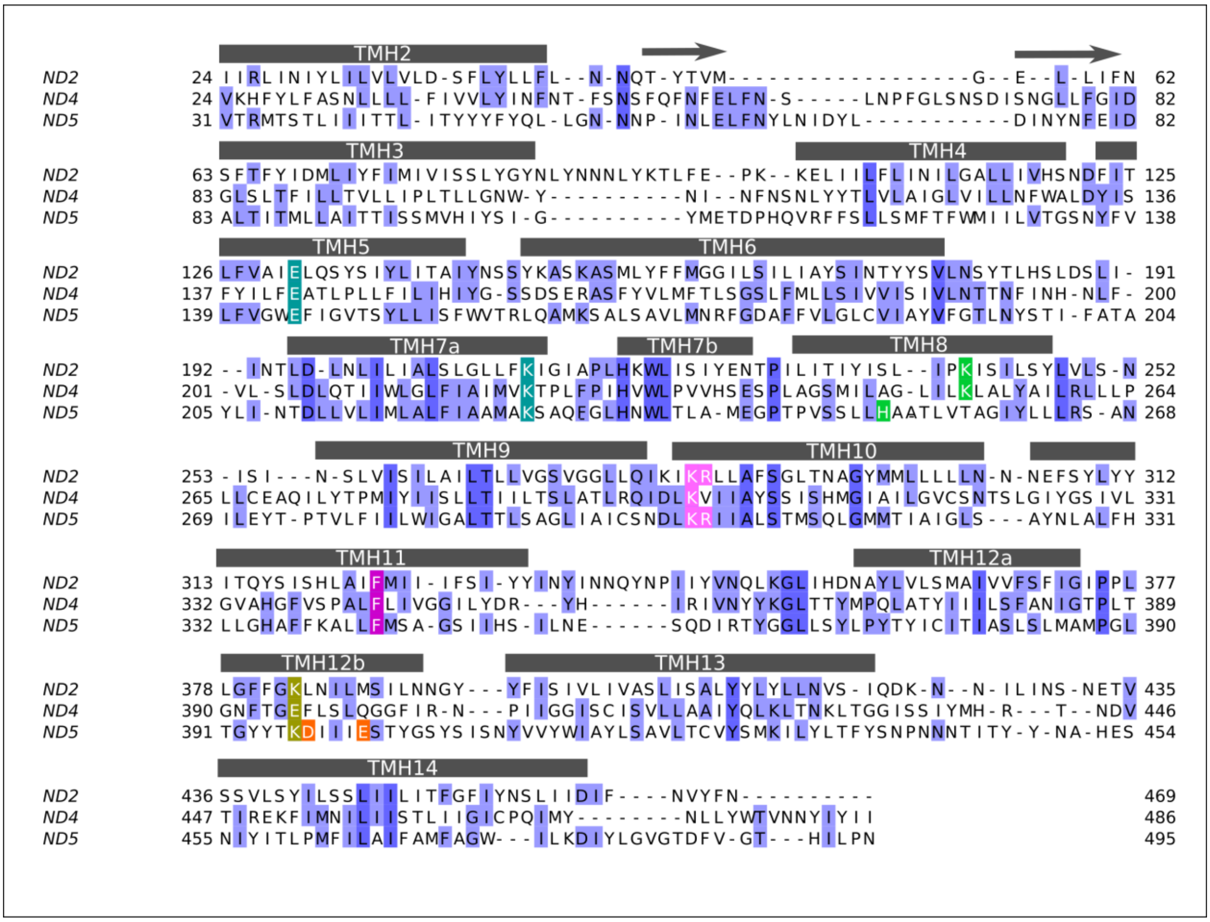


**Figure S4. Multiple sequence alignment of antiporter-like subunits ND2, ND4 and ND5 from *Y. lipolytica***. The TM helices and the β-strands are indicated. The residues highlighted in shades of blue are colored by sequence identity. Amino acid residues discussed in the text are colored; conserved ion-pair from TMH5 and TMH7a (turquoise), conserved Lys/His residues from central hydrophilic axis (green), Lys/Glu from TMH12b (olive green), Asp and Glu from TMH12b (orange) that form proton releasing route in ND5, conserved residues of KR motif, a potential proton uptake site (pink) and conserved Phe from TMH11, a potential "gating" residue (magenta). The TMH1, TMH15 and the lateral helix of ND5 have been removed. Residue Lys511 from TMH15 of ND5 (not shown) is conserved in mitochondrial sequences, and participates in P-side connection of ND5 subunit. Alignment is based on the structural alignment of three subunits, and was prepared using Jalview (Waterhouse et al., 2009).

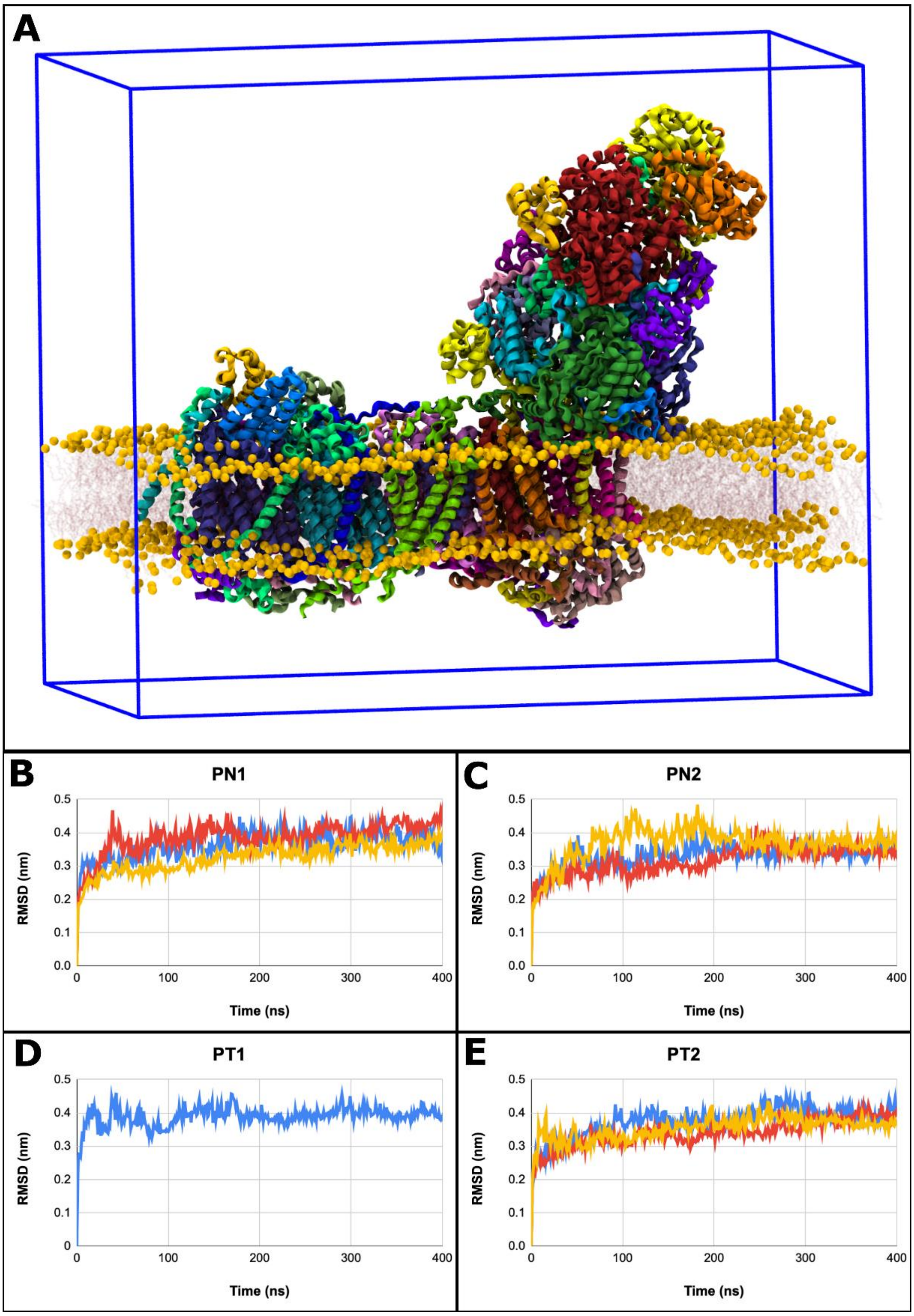


**Figure S5. Molecular dynamics simulation setup and its stability. (A)** Model system of respiratory complex I in membrane-solvent environment. Protein subunits are colored as in main text Figure 1. Lipid head groups are shown as orange spheres and lipid tails are shown in transparent red. The blue box shows the periodic boundaries, which is filled with water and Na and Cl ions (omitted for clarity). **(B-E)** Root mean square deviation (RMSD, in nm) was measured for the protein backbone atoms of all subunits for different simulation setups. The starting configuration (frame 0) of the simulation was used to align the trajectories, and as the reference for the calculation.

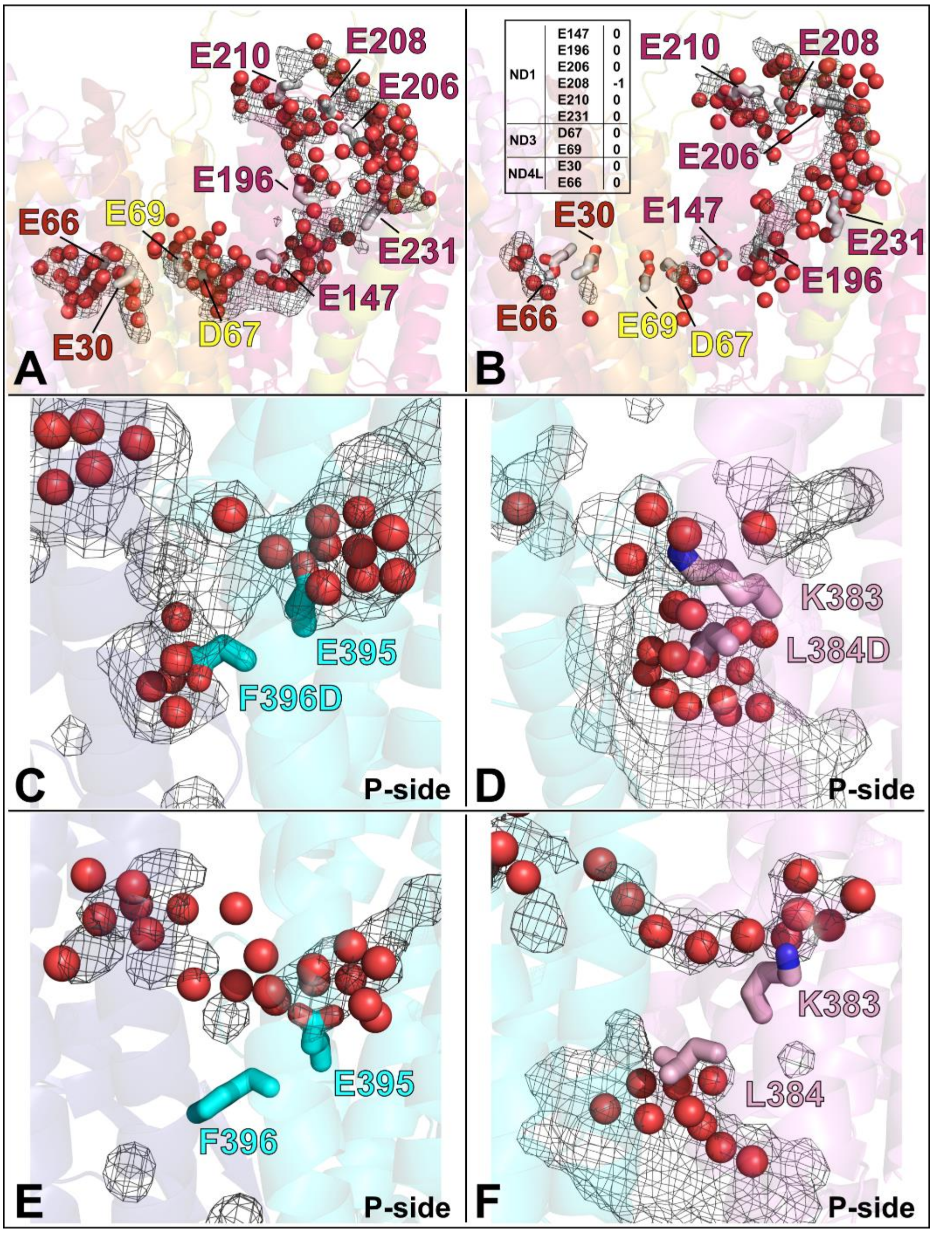


**Figure S6. Hydration in the "E channel" region and induced opening of P-side connections in ND2/4 subunits. (A)** For the E channel region (ND1, hot pink; ND3, yellow; ND4L, red; ND6, orange), the volumetric mesh is shown from MD setup PN1 with all titratable residues in their charged states and **(B)** from MD setup PN2 with protonation states as in the inset table. Water molecules (red spheres) within 6 Å from residues shown in stick representation were selected from entire simulation data. The isovalue of the mesh in panels (A) and (B) is 0.20. **(C)** Effect of introducing F396D (ND4) and **(D)** L384D (ND2) *in-silico* point mutations on the hydration of the P-side regions of antiporter-like subunits. **(E)** Hydrophobic residues Phe396$^{ND4}$ and **(F)** Leu384$^{ND2}$ insulate the protein interior from the aqueous phase at the P side of the membrane in the wild-type enzyme. Panels (C) and (D) are from simulation setup PN3, while (E) and (F) are from PN1. Water molecules are shown within 6 Å of the mutated residues and selected residues shown in Figure 3A. The mesh was calculated by selecting water molecules within 6 Å from these selected residues from the entire simulation data. The isovalue of the mesh is 0.10 in panels C-F. Water molecules are shown in red spheres and residues in sticks using pink (ND2) and cyan (ND4) for carbon atoms, blue for nitrogen and red for oxygen. Subunit coloring follows the same scheme as Figure 3A.

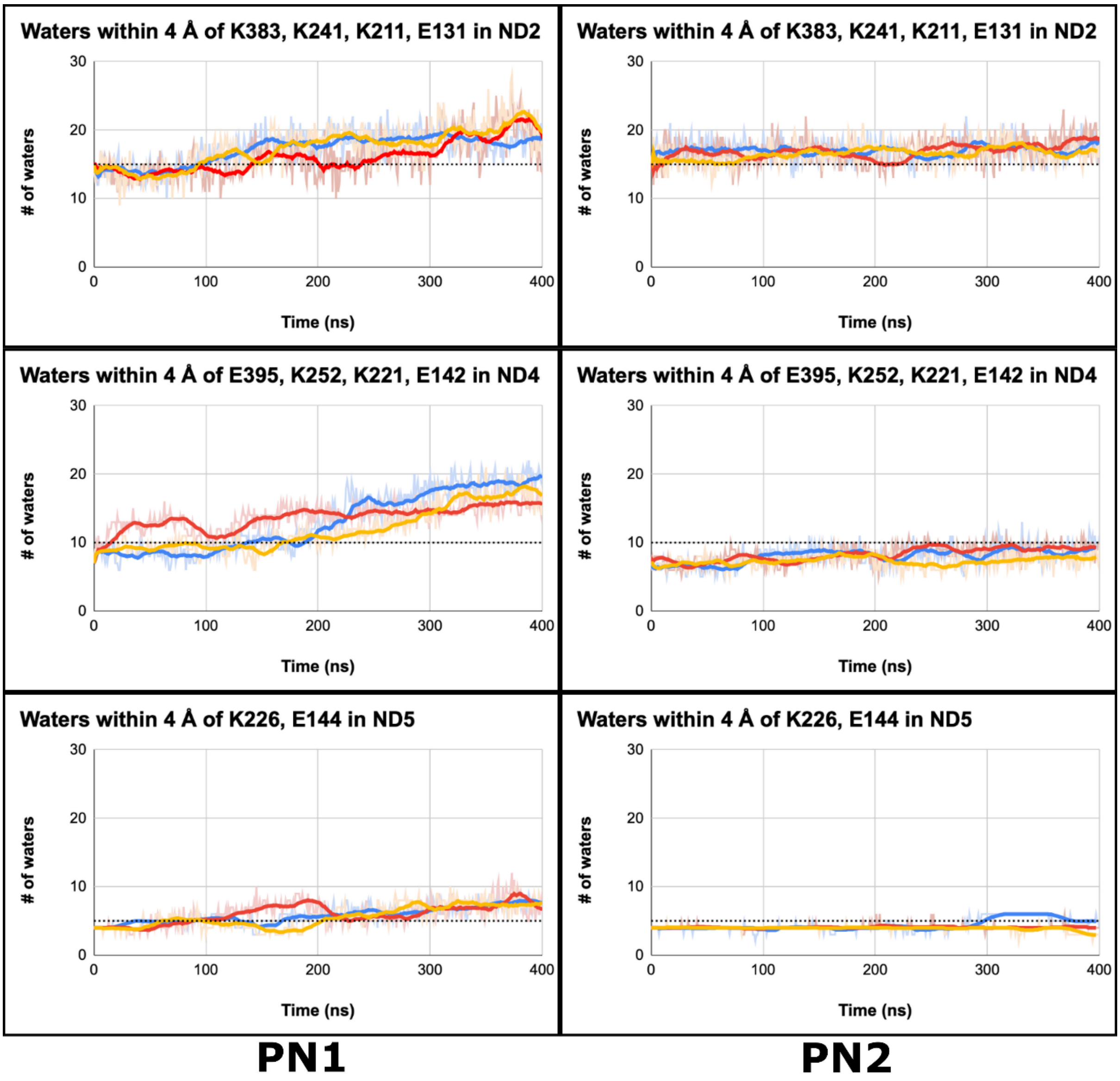


**Figure S7. Water content in central hydrophilic axis of antiporter-like subunits.** The left and right panels show number of water molecules from selected residues in central hydrophilic axis from PN1 and PN2 simulations, respectively. Data from PN2 simulations is closer to structural water content (shown by dotted lines). The three MD replicas are shown for each simulated state in different colors. The bold lines are running average over 20 simulation frames.

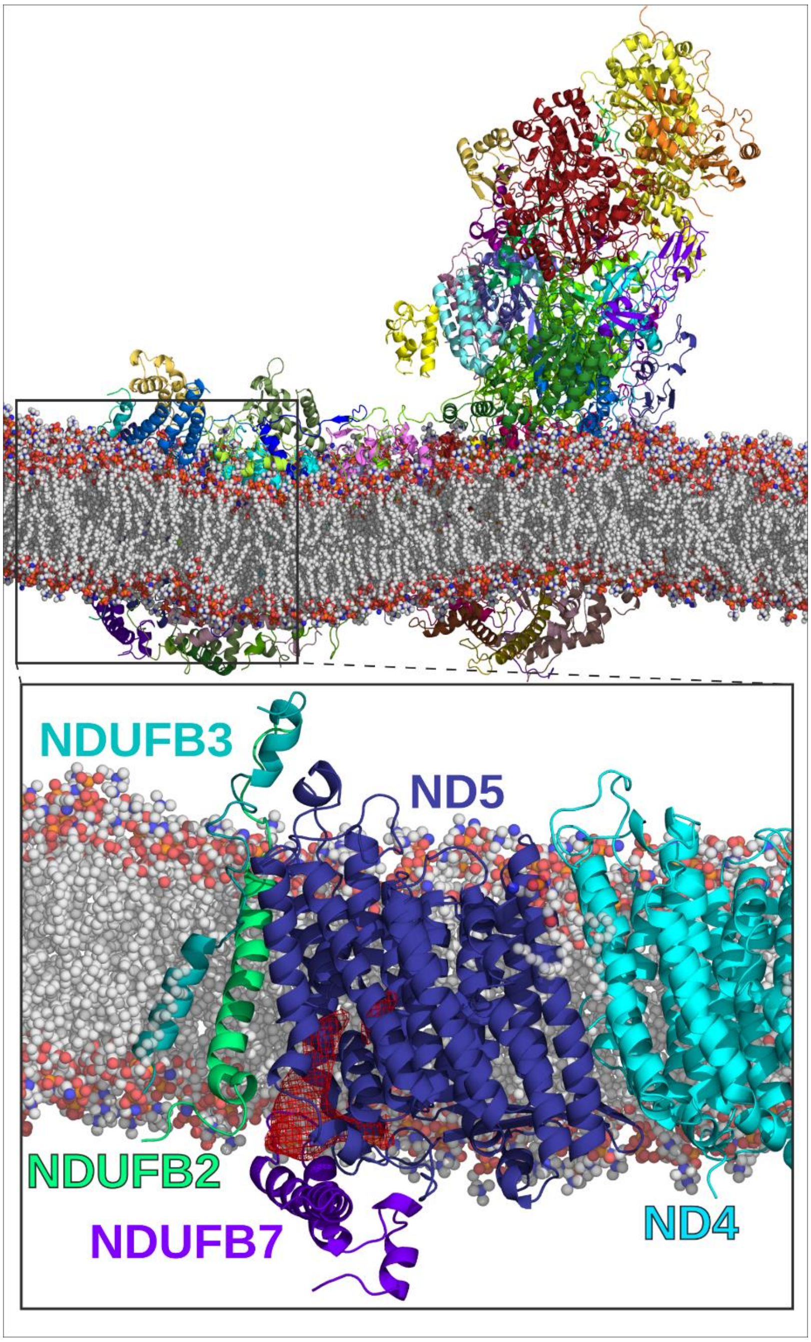


**Figure S8. Membrane bending and lipid-protein interactions.** Lipid bilayer bends and adjusts according to protein structure in MD simulations of complex I. **Inset** shows the unique lipid-protein architecture near putative proton exit route in ND5 subunit, in part induced by tilted TMHs of accessory subunits NDUFB3 and NDUFB2.

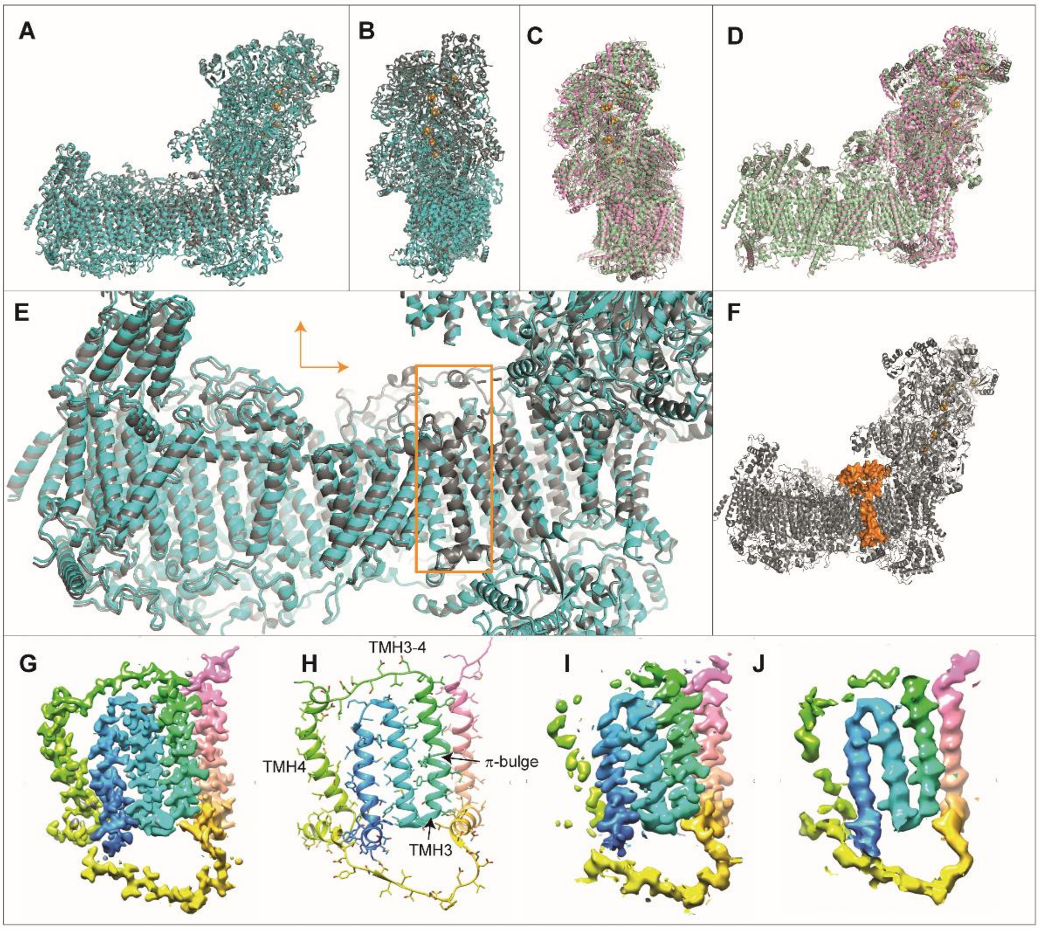


**Figure S9. Overall conformational changes in complex I under turnover. (A)** Sideview of *Y. lipolytica* complex I (deactive, gray; turnover, blue), **(B)** view from the backside, **(C,D)** same views as in (A,B) for an overlay of closed (pink, PDB ID: 6zkc) and open (green, PDB ID: 6zkd) state of ovine complex I (overlayed on subunit ND1), **(E)** same orientation as in (A) showing detailed view of the membrane arm, orange arrows indicate relative movement of $P_D$ module, the orange box encloses an area with weak cryo-EM density for complex I under turnover, **(F)** orange surface highlights protein structure for which cryo-EM density is weak or missing for complex I under turnover, including a section of the long N-terminal extension of NDFUS2, the C-terminus of NDUFA9 and TMH4 and the TMH3-4 loop of ND6, **(G)** segmented map of ND6 from the 2.1 Å complex I map, **(H)** fitted model, **(I)** 3.4 Å map of ND6 under turnover (this work), **(J)** 4.5 Å map, EMD-4385 (Parey et al., 2018); G-J colored from blue (N-terminus) to pink (C-terminus). Under turnover, density for the TM3-4 loop and for TM4 is weak, indicating mobility of these regions, while TM1-3 and TM5 including the π-bulge region of TM3 are rigid.

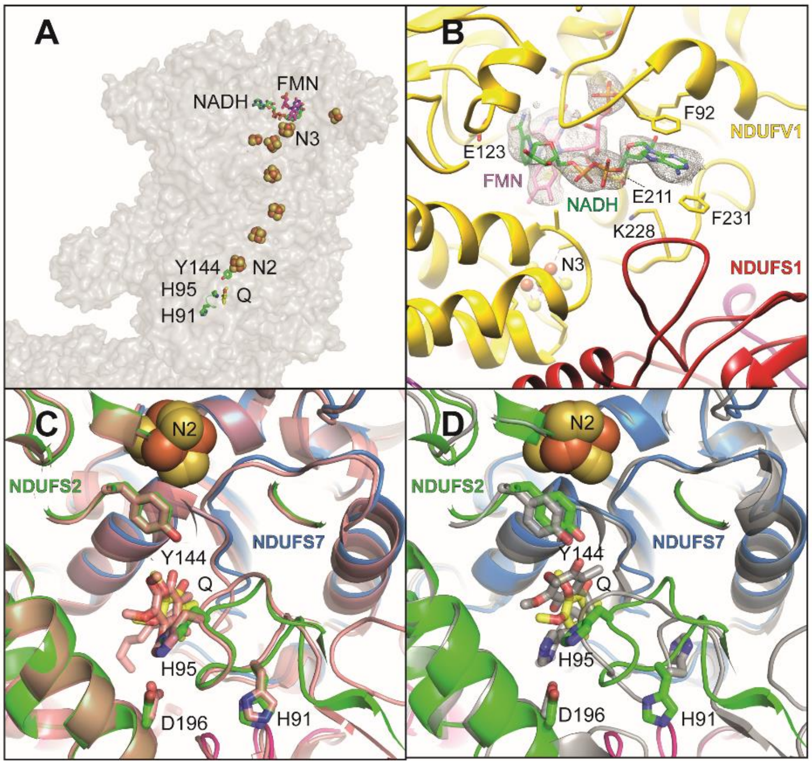


**Figure S10. NADH oxidation and Q reduction site of complex I. (A)** Topology of electron transfer reactions in the peripheral arm, **(B)** density for substrate NADH and the primary electron acceptor FMN in the NADH oxidation site, selected residues participating in NADH binding are shown in stick representation. **(C)** Q reduction site of *Y. lipolytica* complex I captured during steady state activity (DBQ, yellow; compare Figure 4) overlayed with complex I from sheep (salmon, PDB ID: 6zkc) and **(D)** with complex I from *T. thermophilus* (gray, PDB ID: 6i0d). In *Y. lipolytica* and ovine complex I the distance between the tyrosine residue near cluster N2 and Q is too long for a hydrogen bond. A different mode of Q binding is observed in complex I from *T. thermophilus* where Q is bound in a tyrosine – histidine ligation.

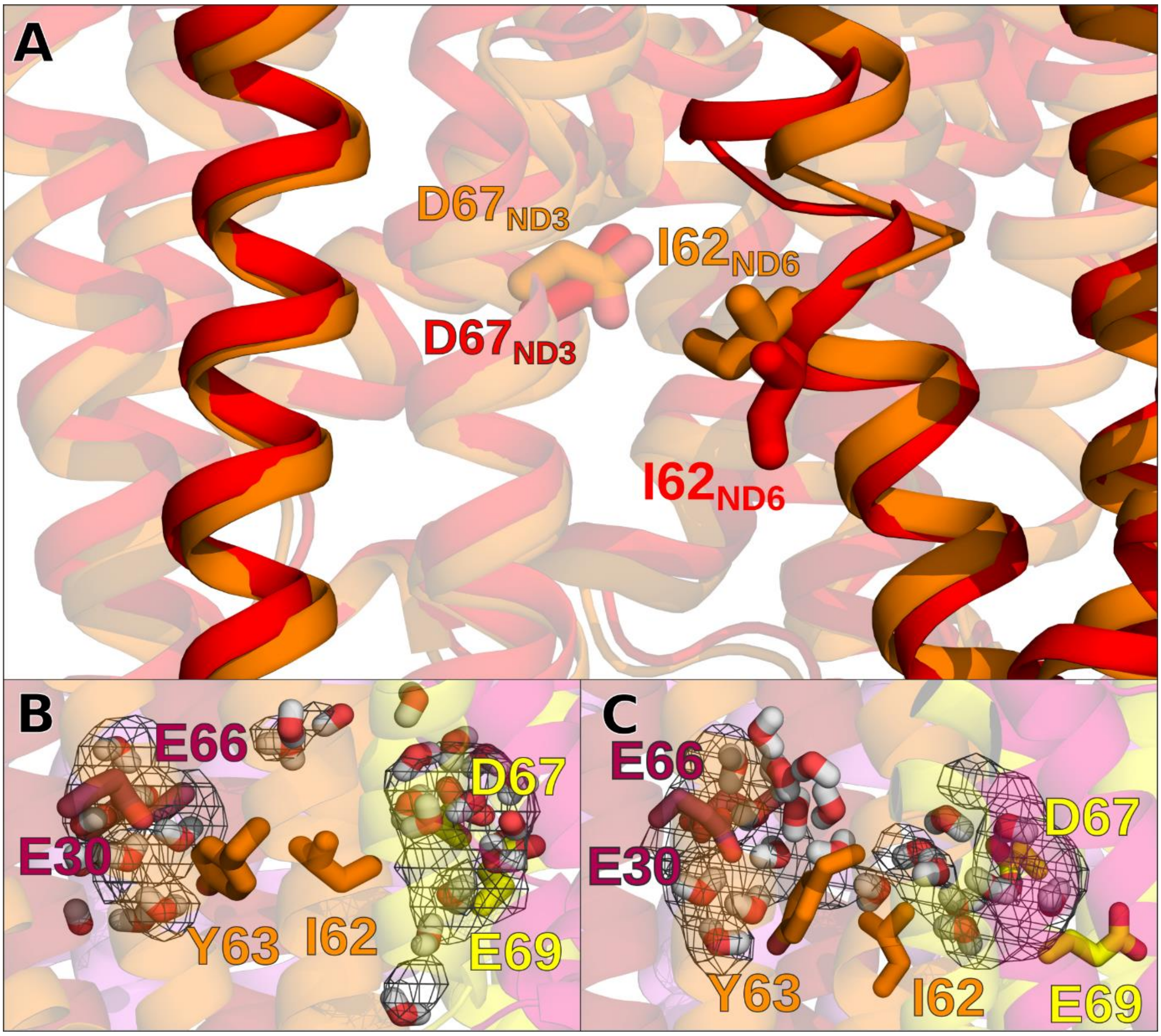


**Figure S11. Hydration bridge between acidic residues of ND3 and ND4L. (A)** Overlay of two MD simulation snapshots showing two conformations of Ile62 of ND6 subunit. Change in conformation of Ile62 and surroundings either closes **(B)** or opens **(C)** the route leading to water-influx in the dry region connecting conserved acidic residues of ND3 and ND4L subunits. The volumetric mesh (0.2 isovalue) is calculated over entire simulation trajectory (setup PN1, simulation replica 3 for panel B and simulation replica 2 for panel C).

**Movie 1.** Cryo-EM density (grey mesh) and model (protein in stick representation; inorganic sulfur, iron and water shown as spheres) close to FeS cluster N2 (colors compare Figure 1).

**Movie 2.** Global conformational changes in respiratory complex I of *Y. lipolytica*. The morph shows the change between complex I in the D form and complex I under turnover conditions. The $P_D$ module slightly moves towards the matrix side and towards the $P_P$ module. There are significant conformational changes at the junction of peripheral arm and membrane arm, most notably in ND1.

**Movie 3.** Conformational changes in ND1 and in the Q binding site. The movie starts with a still image that shows labeling of selected residues. The morph between complex I in the D form and complex I under turnover conditions shows bending of TMH4 of ND1, conformational changes of the ND1 TMH5-6 loop and connected changes in and around the Q binding site. The following short sequence displays the position of the Q molecule in the turnover structure. The movie continues with restoration of the initial state.
